# Single Cell RNA Sequencing and Spatial Profiling Identify Mechanisms of Neonatal Brain Hemorrhage Development and Resolution

**DOI:** 10.1101/2025.07.30.667675

**Authors:** Santiago A. Forero, Zhihua Chen, Ali Pirani, Arpan De, Zachary Wise, John E. Morales, Joseph H. McCarty

## Abstract

Precise control of cell-cell communication networks within brain neurovascular units (NVUs) promotes normal tissue physiology, and dysregulation of these networks can lead to pathologies including intracerebral hemorrhage (ICH). The cellular and molecular mechanisms underlying ICH development and subsequent tissue repair processes remain poorly understood. Here we employed quantitative single cell RNA sequencing coupled with spatial in situ gene expression profiling to characterize NVU signaling pathways associated with ICH in neonatal mouse brain tissue. The initial stages of ICH pathogenesis are characterized by downregulation of extracellular matrix (ECM)-associated signaling factors (Adamtsl2, Htra3, and Lama4) that functionally connect to canonical TGFβ activation and signaling in vascular endothelial cells. Conversely, the progressive resolution of ICH involves upregulation of neuroinflammatory signaling networks (Gas6 and Axl) alongside activation of iron metabolism pathway components (Hmox1, Cp, and Slc40a1) in astrocytes and microglial cells. Integrated computational modeling identifies additional ligand-receptor signaling networks between perivascular glial cells and endothelial cells during both ICH pathogenesis and resolution. Collectively, these findings illuminate the molecular signaling networks that promote NVU maturation and provide novel mechanistic insights into the pathways controlling ICH pathogenesis and repair.

## Introduction

The brain is the most vascularized organ in the mammalian body, with its web of arteries, veins and capillaries interacting with surrounding neurons and glia in multicellular complexes termed neurovascular units (NVUs) ^1^. Growth factors, extracellular matrix (ECM) proteins, and other factors mediate communication between NVU cells to promote normal brain development and physiology ^2^. Perivascular glial cells play especially important roles in promoting NVU assembly and maintaining vascular integrity by activating complex signaling cascades in endothelial cells ^3^. Indeed, during normal brain development, coordinated signaling between perivascular cells and the vascular endothelium induces blood-brain barrier (BBB) formation. Signaling pathways linked to Wnts, vascular endothelial growth factor, and other secreted factors have been identified as crucial in NVU development ^4^. Disruption of these critical signaling pathways can cause BBB defects leading to edema and intracerebral hemorrhage (ICH) that frequently initiate in periventricular neurogenic regions of the developing brain ^5^. Additional pathways in perivascular glial cells and endothelial cells that drive the development of ICH and related neuropathologies remain largely unknown.

The repair of blood vessel defects in ICH pathologies requires coordinated interactions between endothelial cells, pericytes, astrocytes, microglia, and oligodendrocytes, each contributing specialized functions to vascular restoration. Endothelial cells initiate angiogenic responses through microglia-expressed vascular endothelial growth factor ^6^, while pericytes provide structural support and regulate vessel maturation through PDGFB-mediated pathways ^7^. The progressive resolution of ICH also involves pathways that reestablish endothelial cell quiescence and enhance glial stabilization of blood vessels . This is mediated, in part, by the restoration of Notch signaling in endothelial cells, which suppresses proliferative responses and promotes barrier maturation through tight junction proteins ^8^. Astrocytes extend their end feet to reestablish contact with newly formed vessels, secreting factors such as transforming growth factor β (TGFβ) that promote endothelial survival and BBB function ^9^. Astrocytic endfeet reconstitute their polarized distribution of aquaporin-4 and Kir4.1 channels, facilitating proper water homeostasis and potassium buffering around repaired vessels ^10^. While these individual factors have been characterized in ICH repair, their complex temporal and spatial regulation as well as their functional cross talk with components of other critical pathways at the NVU remain unknown.

In this study, single-cell RNA sequencing techniques and in situ spatial transcriptomics platforms have been integrated to generate high-resolution mapping of cellular responses in mouse models of ICH. These efforts have identified previously uncharacterized cell type-specific pathways that are differentially activated during the initiation and resolution phases of hemorrhage. For example, new adhesion and signaling pathways in endothelial cells and astrocytes have been identified that drive ECM remodeling and promote endothelial barrier maturation at the NVU. In addition, we have uncovered microglial-mediated neuroinflammatory pathways involved in restoration of BBB integrity, clearance of iron from extravasated red blood cells, and repair of necrotic neural tissue. Our results bridge fundamental pathways with pathological outcomes, providing critical insights into both normal developmental processes and disease mechanisms at the NVU. This work also advances our understanding of the complex interplay between perivascular glial cells and hemorrhagic blood vessels while identifying potential therapeutic targets for treating patients with ICH.

## Results

### Single-cell transcriptomic profiling of cerebral cortices from mouse models of ICH

Integrin β8/Itgb8 is expressed in perivascular glial cells and plays critical roles in regulating brain NVU formation and BBB maturation via activation of latent-transforming growth factor β (TGFβ) in the ECM and subsequent signaling in endothelial cells ^11–15^. As shown in Fig. 1A, mice with glial cell-specific deficiencies in Itgb8 adhesion and signaling functions display severe cortical ICH at birth (post-natal day 0, P0) that is progressively resolved by P10. The ICH resolution correlates with robust activation of perivascular Iba1^+^ microglia and GFAP^+^ astrocytes (Fig. 1B and Supp. Fig. 1). To identify genes and pathways related to ICH pathogenesis and repair, we performed single cell RNA sequencing (scRNAseq) of cerebral cortices dissected from neonatal Nestin-Cre/+; Itgb8f/f mutant (KO) and Nestin-Cre/+;Itgb8+/+ control pups (WT) at P0, P5, and P10 (Fig. 1C). As a validation of our conditional knockout models using Nestin-Cre ^16^, we detected reduced expression of Itgb8 transcripts in astrocytes from KO samples compared to WT controls. Itgb8 expression is detected in OPCs and OLs in both KO and WT samples at all ages (Supp. Fig. 2), confirming our prior data showing that the Nestin-Cre transgene is active in the neuroepithelial cell lineage that gives rise to many post-natal astrocytes ^17^.

**Figure 1.**
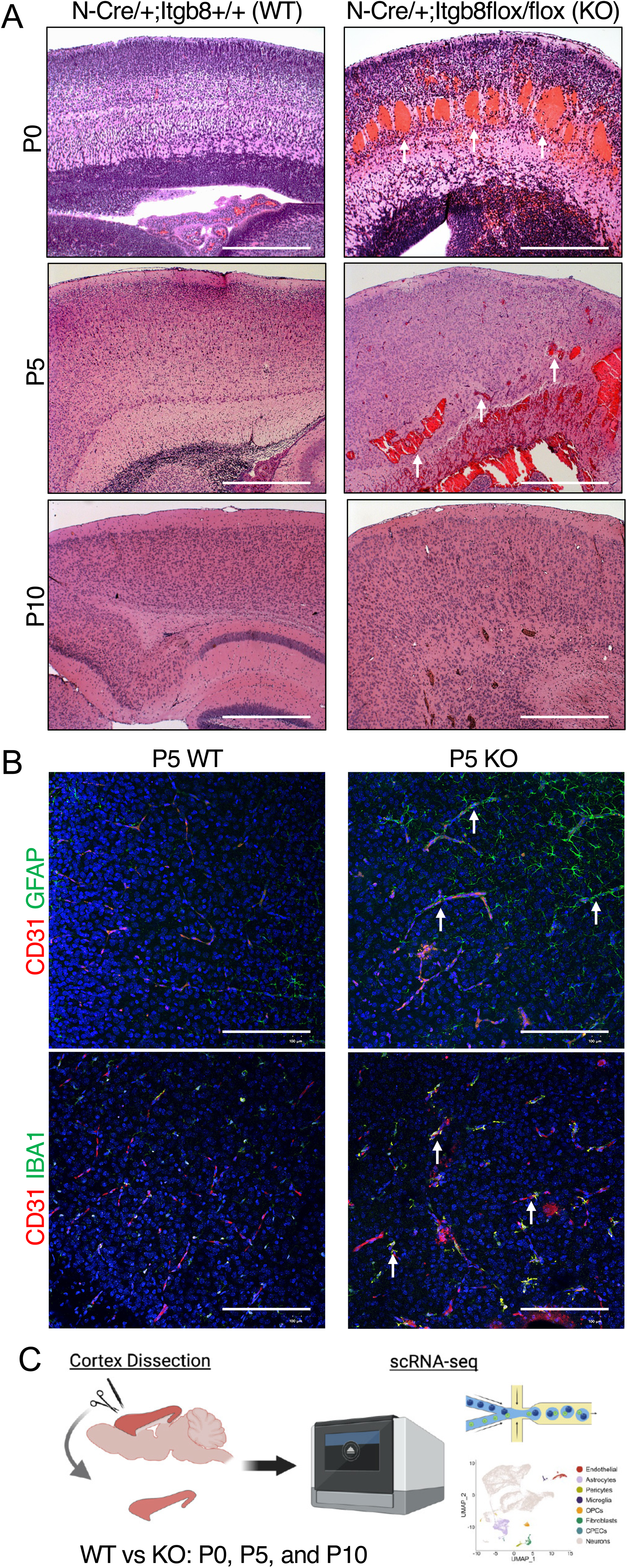
Strategy to analyze mechanisms of ICH pathogenesis and repair by quantitative single cell RNA sequencing. (A); H&E-stained images of sagittal sections through the cerebral cortices of WT (Nestin-Cre;Itgb8 +/+) and KO (Nestin-Cre;Itgb8flox/flox) mice at P0 (upper panels), P5 (middle panels), and P10 (lower panels). Note that KO mice display obvious cortical hemorrhage at P0 and P5 (arrows), which is largely resolved by P10. Scale bars, 50 μm. (**B);** Immunofluorescent analysis of WT (left panels) and KO (right panels) cerebral cortices at P5 using anti-CD31 antibodies to label vascular endothelial cells (red) in combination with anti-GFAP (green) to label astrocytes (top panels) or anti-Iba1 (green) to label microglia (bottom panels). Note the increase in GFAP^+^ and Iba1^+^ reactive astrocytes and microglial cells in the hemorrhagic cortices (arrows). Scale bars, 100 μm. **(C);** Experimental workflow for scRNAseq experiments to identify genes and pathways linked to hemorrhage development and repair in the neonatal cerebral cortex.

Transcriptome profiling revealed comparable numbers and proportions of cell types between KO and WT cortices across all three neonatal ages (Fig. 2A-C). Dimensionality reduction mapping of cell types showed a high degree of overlap in gene expression levels between clusters in KO and WT samples (Fig. 2A-C). Using canonical markers, we identified expected cell types that comprise NVUs: endothelial cells, astrocytes, pericytes, microglia, oligodendrocyte progenitor cells (OPCs)/oligodendrocytes (OLs), fibroblasts and neurons (Fig. 2A-C). As expected, most of the cell clusters identified in the analyses were neuronal populations.

**Figure 2.**
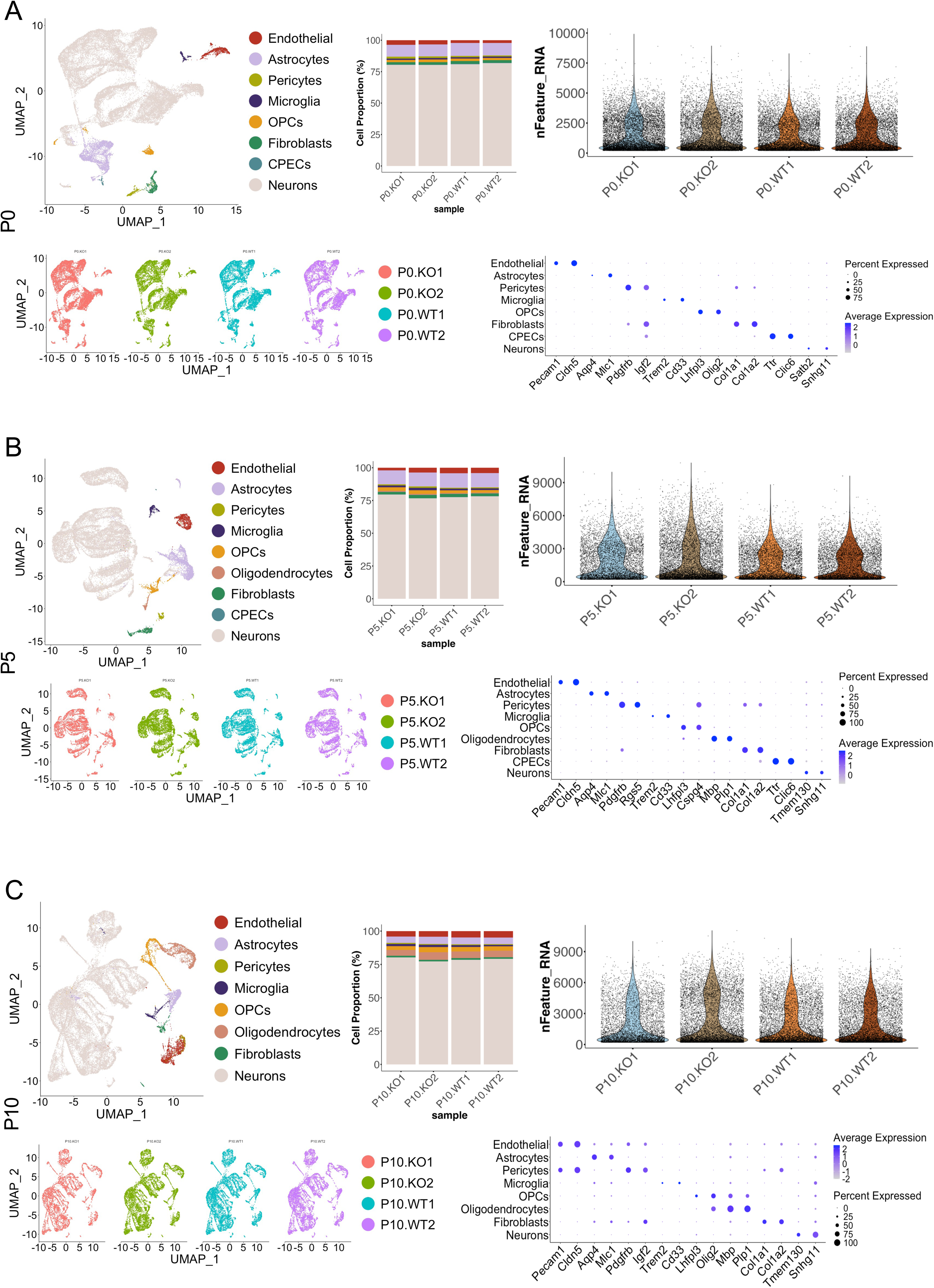
Dimensionality reduction and cluster annotation of scRNAseq data from control and ICH samples. (A); Uniform manifold approximation and projection (UMAP) visualization of integrated WT and KO cells in P0 sample clusters (top left) and UMAPs split by samples (bottom left). Shown are bar plots of cell-type proportions (top middle), violin plots of number of nFeatures (top right), and a dot plot of marker genes used for cluster annotation (bottom right). **(B);** UMAP visualization of integrated P5 control and ICH sample clusters (top left) and UMAPs split by samples (bottom left). Bar plots of cell-type proportions per P5 sample are shown (top middle). Violin plots of number of (top right) and dot plot of marker genes used for cluster annotation (bottom right). **(C);** UMAP visualization of integrated P10 sample clusters (top left) and UMAPs split by samples (bottom left). Bar plots of cell-type proportions (top middle), violin plots of number of nFeatures (top right), and a dot plot of marker genes used for cluster annotation are shown (bottom right).

### Analysis of vascular endothelial cell responses to ICH pathogenesis

We compared ICH-dependent differences in transcript expression for each cell type at the different developmental ages (P0, P5 and P10). At each neonatal age, endothelial cells and microglia from hemorrhagic brains displayed the most changes in gene expression when compared to WT control mice. Differential gene expression analysis revealed that endothelial cells in P0 ICH samples showed a total of 42 upregulated differentially expressed genes (DEGs) and 56 downregulated DEGs (Fig. 3A). Genes upregulated in endothelial cells from ICH samples encode factors involved in endothelial cell adhesion (Basal cell adhesion molecule; Bcam), inflammation (Pleomorphic adenoma gene 1; Plag1), and BBB integrity (Apolipoprotein E; Apoe). Significantly downregulated genes in endothelial cells from hemorrhagic brains were functionally linked to ECM remodeling of basement membranes, such as high-temperature requirement A3 (Htra3), a disintegrin and metalloproteinase with thrombospondin motifs-like 2 (Adamtsl2), Sparc (osteonectin), Cwcv and Kazal-like domains proteoglycan 2 (Spock2), fibronectin (Fn1), laminin-4 (Lama4), and nidogen-1 (Nid1). Gene ontology (GO) enrichment analysis confirmed downregulation of TGFβ-regulated ECM adhesion and signaling pathways in endothelial cells from hemorrhagic cortices (Supp. Fig. 3).

**Figure 3.**
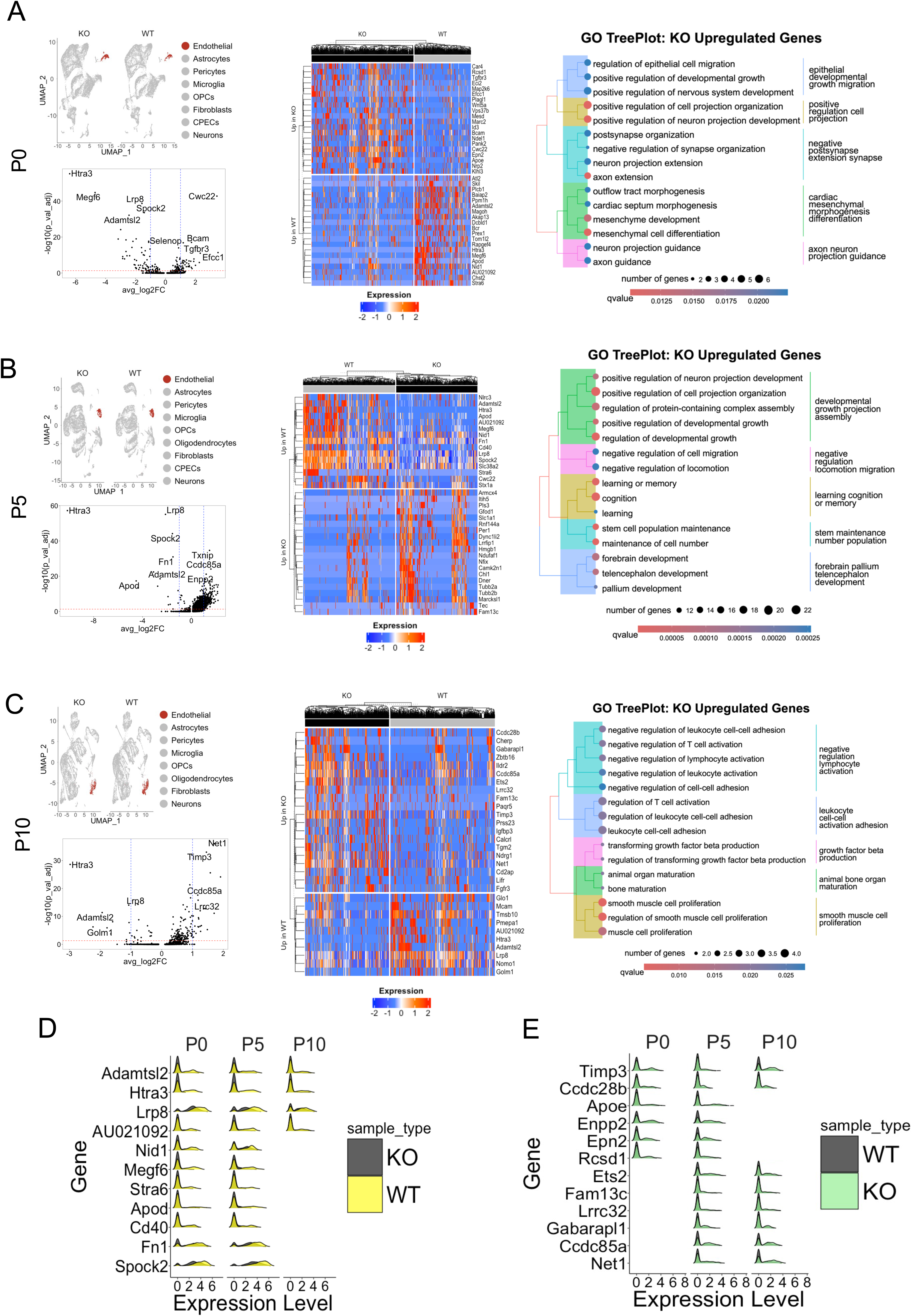
Differential gene expression and GO enrichment analysis of vascular endothelial cells from hemorrhagic cerebral cortices. (A); Condition-split UMAP highlighting endothelial clusters in integrated P0 scRNAseq samples (top left). Volcano plot of significant DEGs in P0 endothelial cells, KO vs. WT (bottom left). Blue lines on volcano plot mark −1 and +1 avg_log2FC cutoff of statistical significance (x axis). Red line on volcano plot marks adjusted p-value significance cutoff (0.05). Y axis is −log10 of adjusted p value. Heatmap showing log-normalized and scaled expression of top 20 DEGs upregulated in P0 ICH samples (up in KO) and downregulated in ICH samples (up in WT). TreePlot of top 15 gene ontology (GO) enrichment pathways overexpressed in endothelial cells at P0 (right). **(B);** Condition-split UMAP highlighting endothelial clusters in integrated P5 scRNAseq samples (top left). Volcano plot of significant DEGs in P5 endothelial cells, KO vs. WT (bottom left). Blue lines on volcano plot mark −1 and +1 avg_log2FC cutoff of statistical significance (x axis). Red line on volcano plot marks adjusted p-value significance cutoff (0.05). Y axis is −log10 of adjusted p value. Heatmap showing log-normalized and scaled expression of top 20 DEGs upregulated in P5 ICH samples (up in KO) and downregulated in endothelial cells from ICH samples (up in WT). TreePlot of top 15 gene ontology (GO) enrichment pathways overexpressed in endothelial cells at P5 (right). **(C);** Condition-split UMAP highlighting endothelial clusters in integrated P10 scRNAseq samples (top left). Volcano plot of significantly differentially expressed genes (DEGs) in P10 endothelial cells, KO vs. WT (bottom left). Blue lines on volcano plot mark −1 and +1 avg_log2FC cutoff of statistical significance (x axis). Red line on volcano plot marks adjusted p-value significance cutoff (0.05). Y axis is −log10 of adjusted p value. Heatmap showing log-normalized and scaled expression of top 20 DEGs upregulated in P10 ICH samples (up in KO) and downregulated in endothelial cells from ICH samples (up in WT). TreePlot of top 15 gene ontology (GO) enrichment pathways overexpressed in endothelial cells from ICH samples at P10 (right). **(D);** Ridgeplot of DEGs downregulated in endothelial cells from KO brains at the three neonatal ages. **(E);** Ridgeplot of DEGs upregulated in endothelial cells from KO brains at P0, P5 and P10.

Endothelial cells analyzed from P5 ICH samples showed 275 upregulated DEGs and 15 downregulated DEGs. Notably, cells displayed upregulation of several genes implicated in BBB integrity (Fig. 3B). Most significantly downregulated genes at P5 overlapped with those identified at P0; for example, Htra3, Megf6, Adamtsl2, Fn1, and Nid1 are downregulated in endothelial cells from P0 and P5 hemorrhagic brains. Furthermore, GO enrichment analysis at P5 revealed downregulation of cell-ECM adhesion and signaling pathways crucial for ECM organization (Supp. Fig. 3).

Endothelial cells from P10 cortical samples showed 24 upregulated DEGs and 10 downregulated DEGs. At this age, the ICH has largely cleared (Fig. 1A). Differential gene expression analysis revealed the enrichment of various genes involved in regulating BBB stability (Fig. 3C). Notably, the tissue inhibitor of metalloproteinases 3 (Timp3), which is linked to inhibiting metalloproteinases that degrade the ECM and adherens junction components ^18,19^, is upregulated in endothelial cells from the hemorrhagic brain. Interestingly, GO enrichment analysis at P10 showed several pathways involved in the regulation of TGFβ signaling (Fig. 3C), further suggesting that at this age, there is significant BBB restructuring and angiogenic activity. Overall, we found six genes in vascular endothelial cells that were consistently expressed differentially in ICH samples at all ages analyzed. Specifically, Adamtsl2, Htra3, LDL receptor-related protein 8 (Lrp8), and AU021092 were downregulated in endothelial cells from hemorrhagic brains at all ages (Fig. 3D) while Timp3 and coiled-coil domain containing 28B (Ccdc28b) were consistently upregulated in endothelial cells from ICH samples (Fig. 3E).

### Analysis of perivascular microglial cell responses to ICH pathogenesis

We next analyzed DEGs in microglia from ICH samples at each neonatal age. P0 microglia from ICH samples showed 52 upregulated DEGs and 41 downregulated DEGs. Several of these DEGs have roles in neuroinflammation, phagocytosis, and tissue repair. In addition to gene signatures of microglial activation, some early signs of likely responses to ICH clearance were detected. For example, we observed an upregulation in expression of solute carrier family 40-member 1 (Slc40a1) (Fig. 4A), which encodes the iron transport protein ferroportin-1 that inhibits cellular ferroptosis due to iron accumulation ^20^. GO analysis validated the enrichment of several endocytosis and phagocytosis pathways (Fig. 4A). Subsequent analysis of microglia from P0 ICH samples revealed genes involved in microglia immune response and neuroinflammation such as sialic acid binding Ig-like lectin-H (Siglech) and hexosaminidase-B (Hexb) ^21–23^. The Slc40a1 gene encodes ferroportin-1 protein which is responsible for iron transport across the BBB and inhibits cellular ferroptosis due to iron accumulation ^20^. Therefore, an upregulation of microglial Slc40a1 in response to hemorrhage presumably plays a role in exporting excess irons they have phagocytosed from leaky red blood cells. Indeed, GO enrichment analysis revealed enrichment of several endocytosis and phagocytosis pathways (Fig. 4A). Subsequent analysis of microglia from P0 revealed genes generally involved in microglia immune response and neuroinflammation such as sialic acid binding Ig-like lectin-H (Siglech) and hexosaminidase-B (Hexb) ^21–23^. GO enrichment analysis identified several pathways related to control of macrophage migration (Fig. 4A, Supp. Fig. 3B).

**Figure 4.**
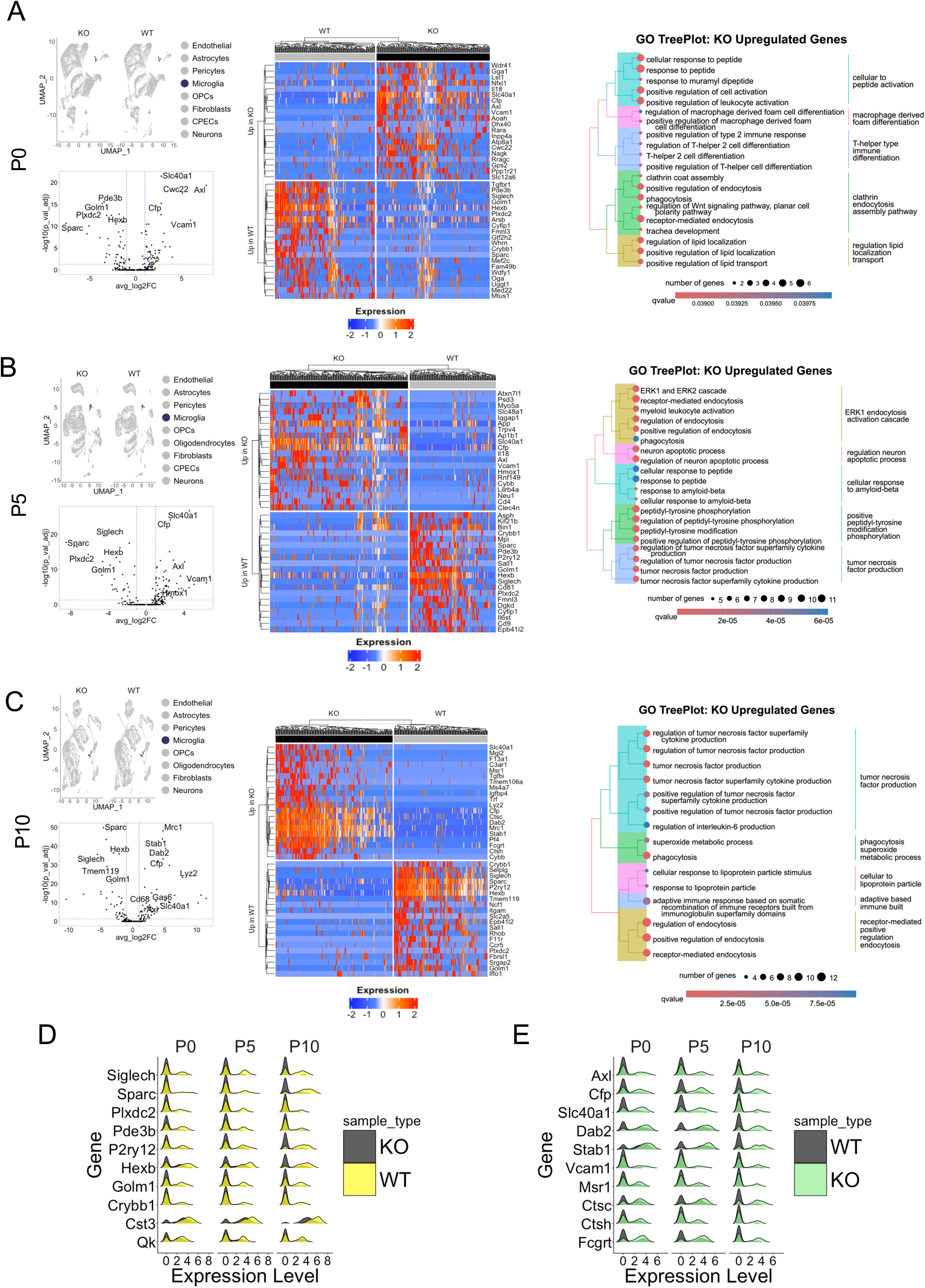
Differential gene expression and GO enrichment analysis of microglia from ICH samples. (A); Condition-split UMAP highlighting microglia clusters in integrated P0 scRNAseq samples (top left). Volcano plot of significantly differentially expressed genes (DEGs) in P0 microglia, KO vs. WT (bottom left). Blue lines on volcano plot mark −1 and +1 avg_log2FC cutoff of statistical significance (x axis). Red line on volcano plot marks adjusted p-value significance cutoff (0.05). Y axis is −log10 of adjusted p value. Heatmap showing log-normalized and scaled expression of top 20 DEGs upregulated in microglia from P0 ICH samples (up in KO) and downregulated in ICH samples (up in WT). TreePlot of top 15 gene ontology (GO) enrichment pathways overexpressed in KO microglia at P0 (right). **(B);** Condition-split UMAP highlighting microglia clusters in integrated P5 scRNAseq samples (top left). Volcano plot of significantly differentially expressed genes (DEGs) in P5 microglia, KO vs. WT (bottom left). Blue lines on volcano plot mark −1 and +1 avg_log2FC cutoff of statistical significance (x axis). Red line on volcano plot marks adjusted p-value significance cutoff (0.05). Y axis is −log10 of adjusted p value. Heatmap showing log-normalized and scaled expression of top 20 DEGs upregulated in microglia from P5 ICH samples (up in KO) and downregulated in ICH samples (up in WT). TreePlot of top 15 gene ontology (GO) enrichment pathways overexpressed in microglia in ICH samples at P5 (right). **(C);** Condition-split UMAP highlighting microglia in integrated P10 scRNAseq samples (top left). Volcano plot of significantly differentially expressed genes (DEGs) in P10 endothelial cells, KO vs. WT (bottom left). Blue lines on volcano plot mark −1 and +1 avg_log2FC cutoff of statistical significance (x axis). Red line on volcano plot marks adjusted p-value significance cutoff (0.05). Y axis is −log10 of adjusted p value. Heatmap showing log-normalized and scaled expression of top 20 DEGs upregulated in P10 ICH samples (up in KO) and downregulated in microglia from ICH samples (up in WT). TreePlot of top 15 gene ontology (GO) enrichment pathways overexpressed in microglia from ICH samples P10 (right). **(D);** Ridgeplot of differently expressed genes downregulated in brain microglia in KO mice at the three neonatal ages (P0, P5, and P10). **(E);** Ridgeplot of shared differently expressed genes upregulated in KO microglia at P0, P5 and P10.

P5 microglial cells isolated from hemorrhagic brains showed 65 upregulated DEGs and 30 downregulated DEGs. Many upregulated DEGs at P5 (Axl, Vcam1 and Il-18) overlap with P0 DEGs (Fig. 4B). Downregulated DEGs at P5, for example Siglech and Hexb, largely overlap with those observed at P0 (Fig. 4B). However, there are several DEGs unique to P5 ICH samples. One such DEG is Hmox1, which encodes the enzyme heme oxygenase-1. Low density lipoprotein receptor-related protein-1 (Lrp1) is also downregulated in microglial from hemorrhagic brains at P5, but not at P0. GO enrichment analysis at P5 reveals the presence of several pathways involved in phagocytosis and macrophage migration. We also detected the presence of gliogenesis and blood vessel remodeling pathways, suggesting early activation of ICH repair mechanisms at P5 (Fig. 4B, Supp. Fig. 3B).

Microglia from P10 ICH samples show 58 upregulated DEGs and 39 downregulated DEGs. Notably, microglia from P10 hemorrhagic cortices express higher levels of Cd68, a key macrophage marker, suggesting these microglia are more phagocytic than at previous ages. We also detected the upregulation of Axl along with its ligand, growth-arrest specific 6 (Gas6).

This receptor-ligand pair has been shown to promote neuroinflammation and phagocytosis ^24^. In addition, we detected upregulation of factor XII (F13a1) and homeostatic iron regulator (Hfe) genes which are responsible for blood coagulation and balancing the iron levels within the brain^25,26,25,26^. GO enrichment analysis show differential regulation of various signaling pathways involved in phagocytosis, endocytosis and superoxide metabolism (Fig.4C). Overall, microglia cells from ICH samples at the three neonatal ages show several conserved DEGs, suggesting a critical response to ICH . Consistently downregulated DEGs include Siglech, Sparc, Plexin domain containing 2 (Plxdc2), phosphodiesterase-3B (Pde3b), Purinergic receptor P2Y, G-protein coupled 12 (P2ry12), Hexb, Golgi membrane protein-1 (Golm1), Crystallin, beta B1 (Crybb1), Cystatin C (Cst3), and Quaking, KH domain containing RNA binding (Qk) (Fig. 4D).

These data reveal a gene signature of abnormal microglial homeostasis and inability to regulate stress and neuroinflammation. Consistently upregulated DEGs (Axl, Complement factor properdin (Cfp), Slc40a1, Disabled 2 (Dab2), Stabilin-1 (Stab1), Vcam1, Macrophage scavenger receptor-1 (Msr1), Cathepsin-C (Ctsc), Cathepsin-H (Ctsh), and Fc fragment of IgG receptor and transporter (Fcgrt) show a signature of the reactive, phagocytic immune response (Fig. 4E). Axl is upregulated in microglia from the hemorrhagic brain at each neonatal age studied.

Additionally, a signature of iron handling and oxidative stress response is evident with Slc40a1. Furthermore, the sustained overexpression of Vcam1, suggests immune cell or vascular interactions of microglia in response to ICH.

### Single cell spatial characterization of endothelial and microglial DEGs

We next combined a custom 100 gene probe panel designed from the scRNAseq data with a 10X Genomics Xenium 5,000 gene panel to profile the spatial expression of transcripts involved in hemorrhage development and clearance (Figs. 2-4). A total of 3 KO and 3 WT brains (one for each neonatal age; P0, P5, P10) were analyzed using the Xenium workflow (Fig. 5A). Spatial analysis showed different cell spatial positions in ICH samples versus control brain samples. Dimensionality reduction of the Xenium samples revealed the presence of multiple cell clusters (endothelial cells, astrocytes, pericytes, microglia, OPCs/OLs, fibroblasts, and neurons), across the three developmental ages with neurons being the predominant cell type (Fig. 5B-D).

**Figure 5.**
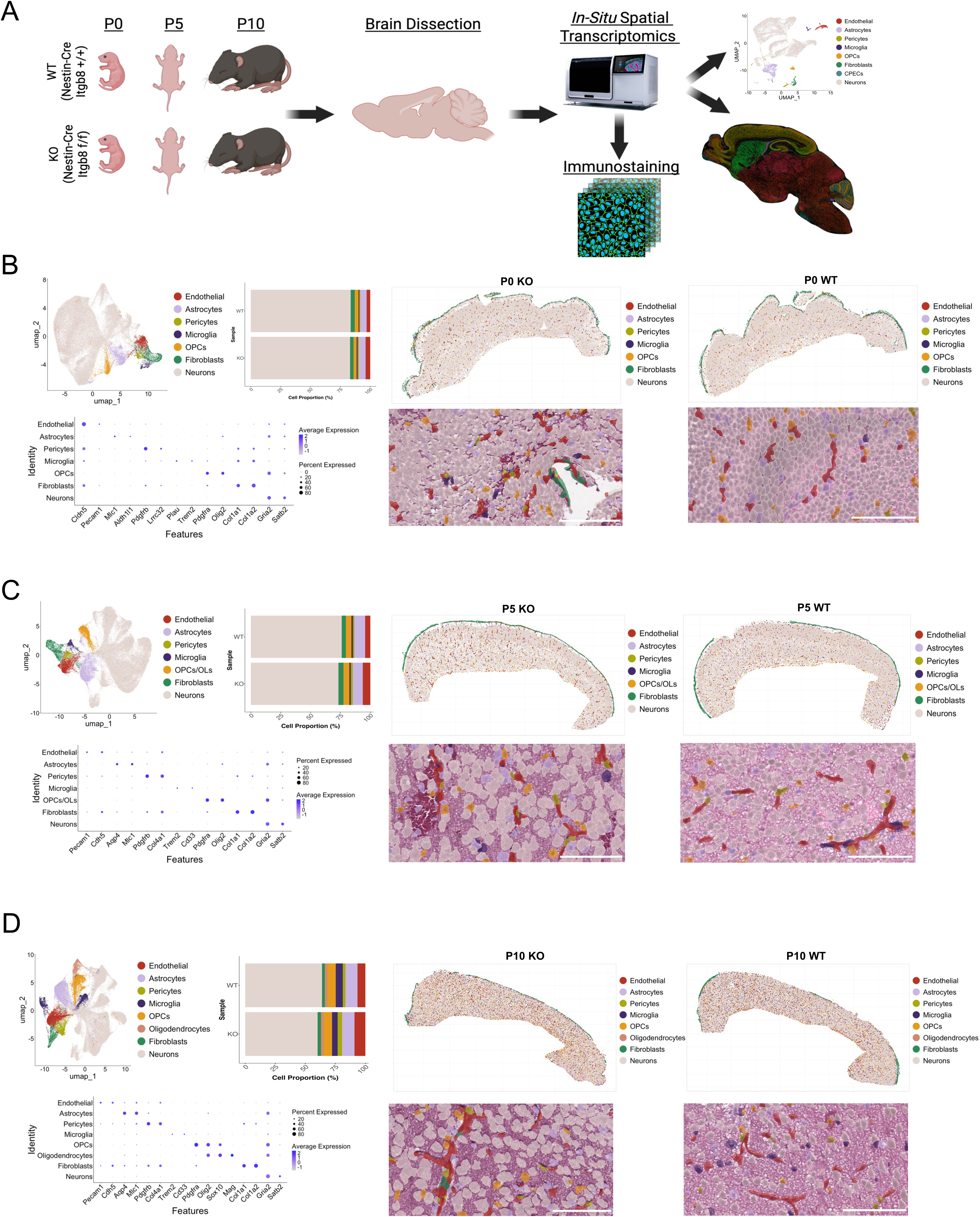
In situ spatial analysis of DEGs in control and ICH samples. (A); Experimental workflow for Xenium-based analysis of gene expression. One mouse was used per age, for each condition. **(B);** UMAP of P0 integrated WT and KO samples (left). Bar plots of P0 cell type proportion per condition (middle top). Dot plot of P0 marker genes used for cluster annotation (bottom left). Spatial location of cells in WT and KO P0 mouse cortex (top right) and H&E images of control and hemorrhagic brains with pseudocolored cells (bottom left). **(C);** UMAP of P5 integrated ICH and control samples (left). Bar plot of P5 cell type proportion per condition (middle top). Dot plot of P5 marker genes used for cluster annotation (top right). Spatial location of cells in WT and KO P5 mouse cortex (top right) and H&E images of control and ICH samples with pseudocolored cells (bottom left). **(D);** UMAP of P10 integrated control and hemorrhagic brain samples (left). Bar plot of P10 cell type proportions per condition (middle top). Dot plot of P10 marker genes used for cluster annotation (top right). Spatial location of cells in WT and KO P10 mouse cortex (top right) and H&E images of samples with pseudocolored cells (bottom left).

In concurrence with the scRNAseq data (Figs. 2-4), spatial analysis of endothelial cells and microglia from hemorrhagic brains show similar DEGs at P0, P5 and P10. At P0, endothelial cells show an upregulation of caveolin-1 (Cav1), and microglia show an upregulation Hmox1. Spatial analysis revealed these transcripts are expressed in microglia around pathological blood vessels (Figs. 6B, 7A). P5 endothelial and microglia DEGs identified by spatial profiling validate those DEGs identified from the scRNAseq analyses. A significant decrease in expression of Htra3 in endothelial cells and Siglech in perivascular microglia is seen directly in hemorrhagic regions (Figs. 6C, 7B). The P10 spatial data also replicate results from the scRNAseq experiments. Adamtsl2 in endothelial cells is significantly downregulated in ICH samples, whereas Axl in microglia is upregulated (Figs. 6D, 7C). Spatial analysis clearly showed this transcript enrichment in perivascular cells in ICH samples. At all ages, cell types showed a high degree of overlap, except for microglia at P10 which showed two distinct clusters in control and ICH samples (Fig. 7C). Although this cluster separation was not present in the microglia from the scRNAseq analysis, differences in DEGs in hemorrhagic brains suggest microglial activation signatures. In particular, by single cell heat map analysis an upregulation of Cd68 and a downregulation of Tmem119 was detected in ICH samples, which is indicative of microglial activation (Fig. 7C)^27,28^.

**Figure 6.**
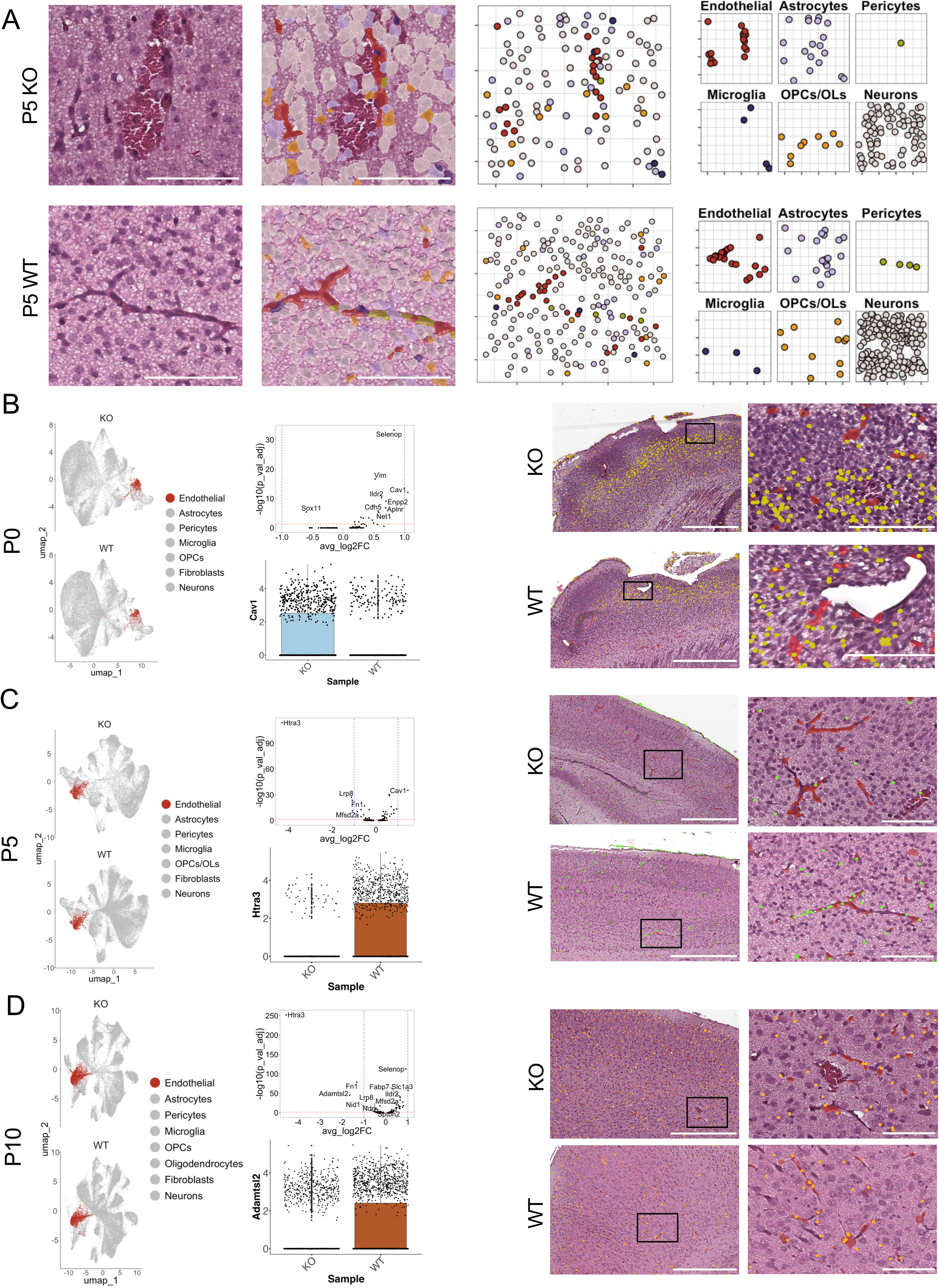
Spatial analysis of endothelial cell DEGs. (A); Representative image of spatial organization of cells around blood vessels in P5 control and ICH samples. **(B);** P0 integrated split UMAP highlighting endothelial clusters (left). Volcano plot of P0 DEGs in endothelial cells from control and ICH samples (top middle). Boxplots of Cav1 expression in P0 endothelial cells (bottom middle). Representative spatial images from control and hemorrhagic cortices showing H&E with overlayed endothelial cells (red) at 500 and 100 μm. Cav1 transcript counts are shown within cells (yellow). **(C);** P5 integrated split UMAP highlighting endothelial clusters (left). Volcano plot of P5 DEGs for endothelial cells from WT and KO brain samples (top middle). Boxplots of Htra3 expression in P5 endothelial cells (bottom middle). Spatial representative images from P0 control and ICH cortex showing H&E with overlayed endothelial cells (red) at 500 and 100 μm. Htra3 transcript counts are shown within cells (green). **(D);** P10 integrated split UMAP highlighting endothelial clusters (left). Volcano plot of P10 DEGs for endothelial cells from WT and KO brains (top middle). Boxplots of Adamtsl2 expression in P0 endothelial cells (bottom middle). Spatial representative images from P10 WT and KO cortex showing H&E with overlayed endothelial cells (red) at 500 and 100 μm. Adamtsl2 transcript counts are shown within cells (orange).

**Figure 7.**
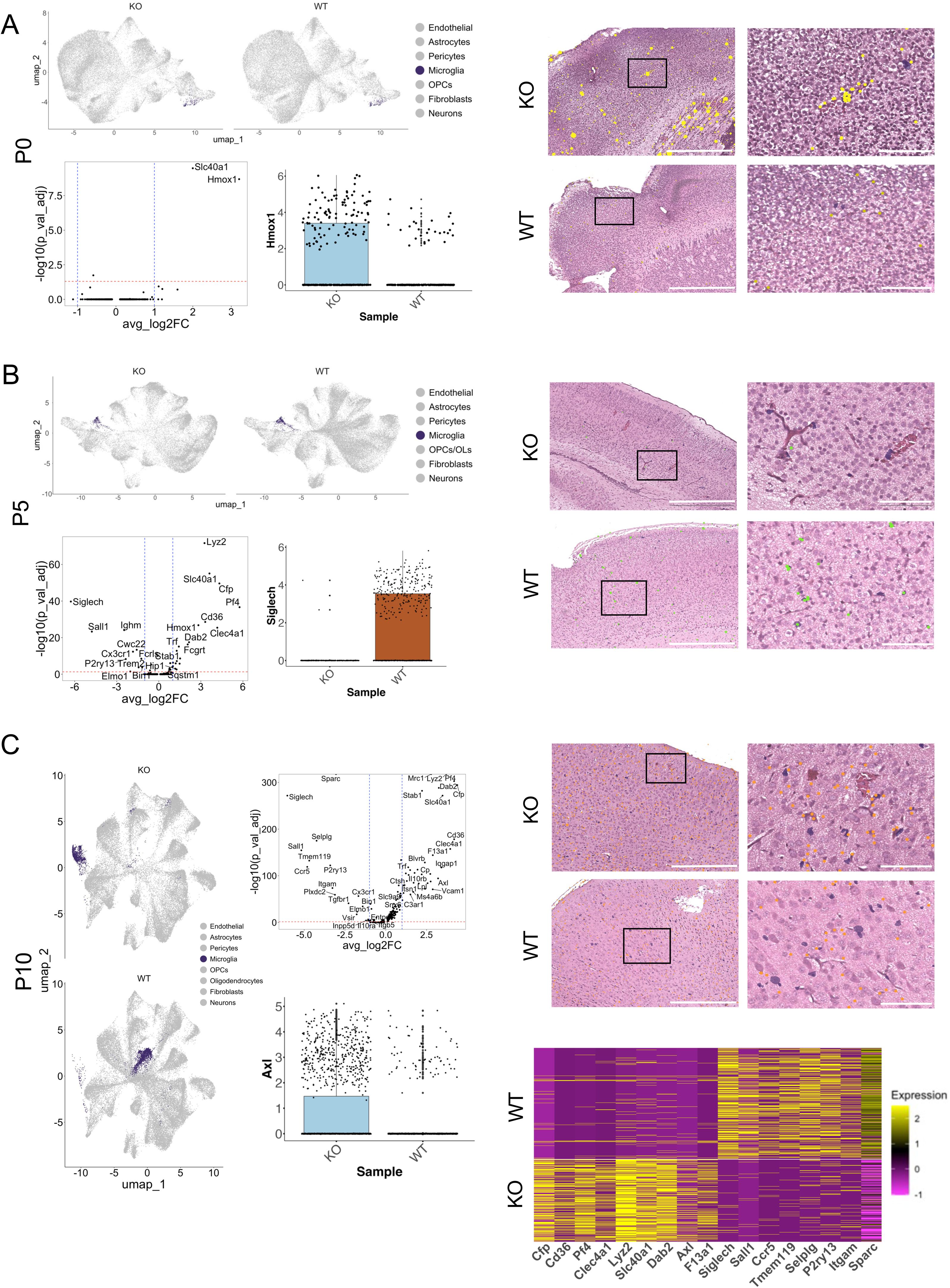
Spatial analysis of microglial cell DEGs. (A); P0 integrated split UMAP highlighting microglia clusters (top left). Volcano plot of P0 DEGs for microglia in WT and KO brains (bottom left). Boxplots of Hmox1 expression in P0 microglia (bottom middle). Spatial representative images (right) from P0 WT and KO cortex showing H&E overlayed microglia (red) at 500 and 100 μm. Hmox1 transcript counts are shown within cells (yellow). **(B);** P5 integrated split UMAP highlighting microglia clusters (left). Volcano plot of P5 DEGs in microglia from WT and KO brain samples (top middle). Boxplots of Siglech expression in P5 microglia (bottom middle). Spatial representative images (right) from P5 WT and KO cortex showing H&E and overlayed microglia (red) at 500 and 100 μm. Siglech transcript counts from differential gene expression analysis are shown within cells (green). **(C);** P10 integrated split UMAP highlighting microglia clusters (left). Volcano plot of P10 DEGs in microglia from WT and KO brains (middle top). Boxplots of Axl expression in P10 microglia (middle bottom). Spatial representative images (top right) from P0 WT and KO cortex showing H&E with overlayed microglia (red) at 500 and 100 μm. Axl transcript counts are shown within cells (orange). Single cell heatmap of scaled transcripts highlighting top DEGs in the two P10 WT and KO microglial clusters (bottom right).

### Pseudotime trajectory inference of gene expression across developmental ages

To analyze ICH-dependent dynamic temporal gene expression patterns, we implemented a pseudotime trajectory analysis of control and hemorrhagic samples (P0, P5, and P10). Merged dimensionality reduction of endothelial cells from all our samples revealed 13 clusters (Fig. 8A), with distinct ages clustering together (Fig. 8B). Slingshot trajectory analysis revealed unique developmental trajectories resulting from hemorrhage (Fig. 8C), suggesting that endothelial cells in ICH samples undergo a distinct differentiation trajectory. Analysis of clusters along the ICH-dependent trajectory revealed unique signatures of gene expression at each age (Fig. 8D). GO enrichment analysis of DEGs revealed the presence of signaling pathways involved in the response to hemorrhage development and resolution. At P0, the major signaling pathways are involved in cellular transport, cellular organization and cell health (Fig. 8E). Notably, signaling pathways like “cell-matrix adhesion”, “cellular component disassembly”, and “regulation of autophagy” stood out as potential mechanisms of cellular re-organization in response to ICH. Analysis of endothelial cells at P5 revealed the presence of “response to iron ion”, “cellular response to metal ion”, and “response to hypoxia” within the top signaling pathways (Fig. 8E, F). The presence of these pathways along with additional apoptotic and catabolic pathway signatures suggests that at P5 (more than at other ages), endothelial cells are responding to hemorrhage. Indeed, the presence of the signaling pathways “blood vessel remodeling”, “wound healing”, and “endothelial cell differentiation” (Fig. 8E, F) suggesting that ICH repair is ongoing, leading toward amelioration of ICH at P10 (Fig.1B).

**Figure 8.**
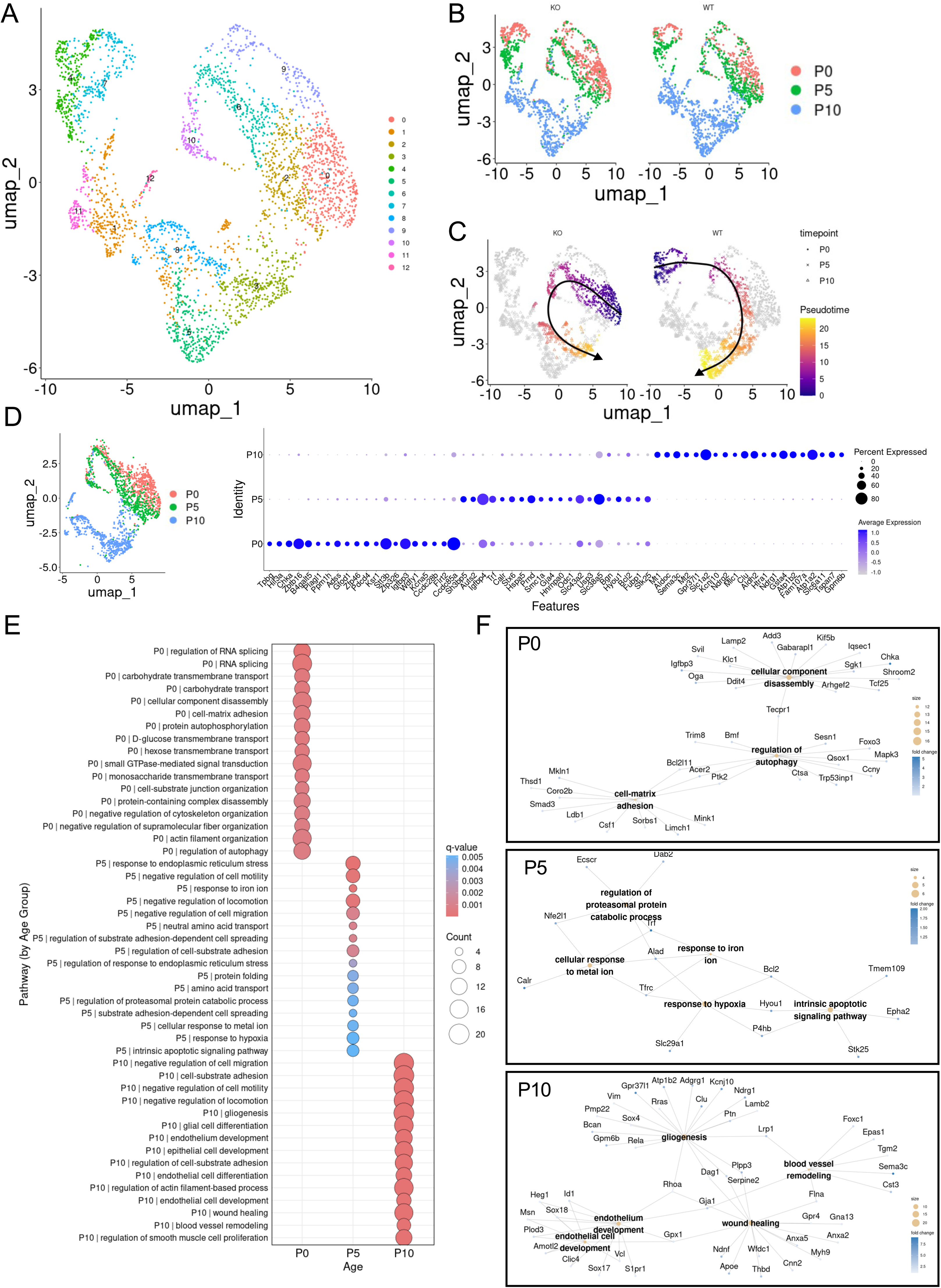
Pseudotime trajectory inference analysis, differential gene expression, and gene ontology enrichment in vascular endothelial cells. (A); UMAP of endothelial cells from all scRNAseq samples. **(B);** UMAP split by condition and clustered according to neonatal age (P0, P5 and P10). **(C);** Condition-split UMAP showing cells within pseudotime trajectory highlighted, cell colors correspond to time along trajectory. Principal curves are shown as black line along trajectory. Arrows show pseudotime trajectory directionality. **(D);** Subset UMAP showing only cells within pseudotime trajectory (colored cells in C), clustered by sample age (left). Dot plot of top 20 DEGs for each age from cells within pseudotime trajectory (right). **(E);** Dot plot of GO enrichment results showing top 15 enriched pathways per neonatal age. **(F);** Key pathways of interest depicted as a category network plot (cnetPlot) showing GO terms and related genes for each neonatal age.

Merged dimensionality reduction of microglial cells from control and ICH samples revealed 10 distinct clusters (Fig. 9A). There are more age-mixed clusters for microglial cells (Fig. 9B) compared to endothelial cells (Fig. 8B). Slingshot trajectory analysis again revealed different developmental paths (Fig. 9C), revealing that microglial cells in the hemorrhagic brain undergo a distinct differentiation trajectory. Analysis of microglial DEGs along the developmental trajectory showed unique gene signatures at each age (Fig. 9D). GO enrichment analysis identified various catabolic pathways (endocytosis, phagocytosis, macroautophagy) active in P0 and P5 brains while P10 microglia cells show DEGs with links to cellular development pathways-synapse organization and junction assembly (Fig. 9E, F). These GO enrichment results suggest that most of the microglial activity in response to ICH clearance and repair is occurring during the early neonatal ages. As mentioned previously, Hmox1 plays a major role in catalyzing the degradation of heme molecules present after serious brain injury ^29^. Hmox1 transcript expression was highly enriched in microglia adjacent to hemorrhagic blood vessels (Fig. 7A). Furthermore, anti-Hmox1 immunostaining confirmed that this increased level of expression was also present at the protein level (Supp. Fig. 4).

**Figure 9.**
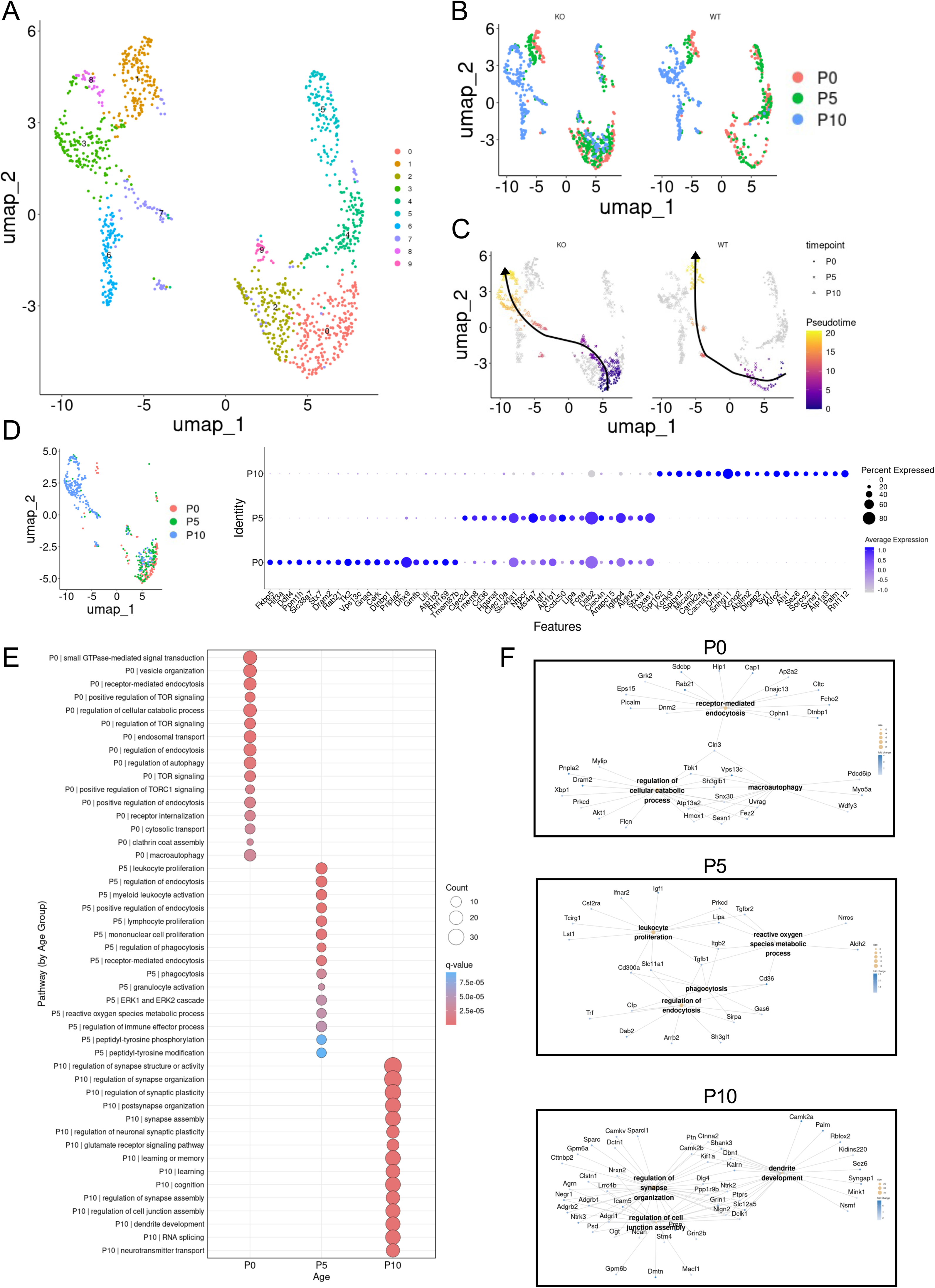
Pseudotime trajectory inference analysis, differential gene expression, and gene ontology enrichment in microglial cells. (A); UMAP of microglial cells from all scRNAseq samples. **(B);** UMAP split by condition and clustered according to neonatal age of sample. **(C);** Condition-split UMAP showing cells within pseudotime trajectory highlighted, cell colors correspond to time along trajectory. Principal curves are shown as black line along trajectory. Arrows show pseudotime trajectory directionality. **(D);** Subset UMAP showing only cells within pseudotime trajectory (colored cells in C), clustered by sample age (left). Dot plot of top 20 DEGs for each neonatal age from cells within pseudotime trajectory (right). **(E);** Dot plot of GO enrichment results showing top 15 enriched pathways per developmental age. **(F);** Key pathways of interest depicted as a category network plot (cnetPlot) showing GO terms and related genes for each neonatal age.

### Computational analysis of Itgb8-dependent cell-cell communication

We next used computational strategies to analyze integrin-dependent communication networks between different cortical cell types at P0, P5 and P10. CellChat v2.1.0 was used to analyze cell-cell interactions within targeted NVU regions across control and ICH samples (Fig. 10A and Supp. Fig. 5). The analysis was conducted in a contact- or spatial distance-dependent manner, utilizing the truncated mean method to compute average gene expression and communication probabilities of signaling pathways using CellChatDB.mouse database. Differential interactions between cell types were identified using the Wilcoxon rank-sum test. LAMA4 signaling was progressively increased in control brain samples at P5 and P10, mainly from endothelial cells and OPCs to astrocytes and endothelial cells (Fig. 10B). In contrast, ICH samples showed elevated LAMA4 signaling early at P0, primarily in endothelial cells, pericytes, and fibroblasts. GAS6 signaling was specifically enriched at P10 in ICH samples. Increased activity of GAS6-AXL and GAS6-MERTK interactions was observed in microglia and fibroblast-to-microglia communication (Fig. 10C). In comparison, WT samples showed minimal GAS6 signaling at this time point. Analyzing signaling interactions specifically between endothelial cells, astrocytes, and microglia showed Galectin, Wnt, and Cxcl pathway activation between endothelial cells and microglia or astrocytes in ICH samples (Fig. 10B). Differential analysis of endothelial cells showed increased Laminin signaling activities at P0 timepoint in KO specifically and decreased FN1 signaling at P10. Astrocytes showed elevated BMP pathway signaling at P10, whereas microglia showed enriched Galectin and Laminin pathway activities at P10 (Supp. Fig. 6), indicating roles for Gas6 signaling during the ICH resolution process.

**Figure 10.**
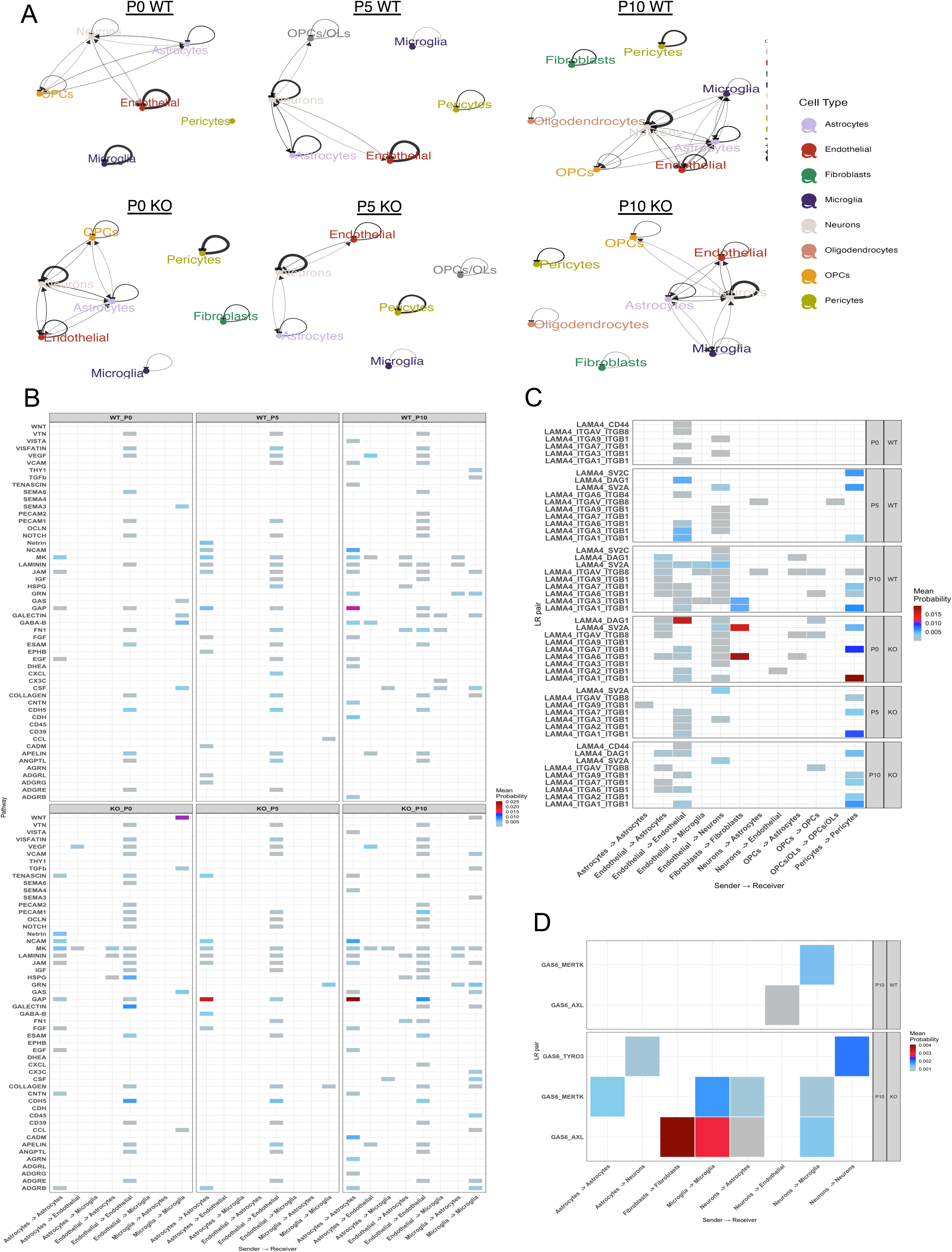
CellChat-based identification of cell-cell communication networks. (A); Chord plot showing direction and interaction weight indicated by arrows/thickness representing aggregated cell-cell communication probability computed using the CellChat probabilistic model. (**B**); Heatmap of communication probabilities between sender and receiver cell types (x-axis) mediated by ligand–receptor pathways (y-axis) for P0, P5, and P10 neonatal ages. Communication probabilities were calculated based on the presence of ligand-receptor factors within cells from the Xenium spatial data. Rectangle color corresponds to mean probability score. **(C);** Heatmap showing examples of communication probabilities across ligand-receptor pairs within the LAMA4 signaling pathway. Sender–receiver cell type interactions (x-axis) for control and ICH samples are shown at the three neonatal ages. Rectangle color corresponds to mean probability score. **(D);** Heatmap of cell-cell communication probabilities between sender and receiver cell types (x-axis) mediated by ligand–receptor pairs involved in GAS signaling pathways (y-axis) for P10 control and ICH samples. Rectangle color corresponds to mean probability score.

## Discussion

In this study, we have identified a network of pathways involved in both the pathogenesis and repair of hemorrhagic brain insults, including several genes with links to ECM stability, BBB integrity, and glial and immune responses . For example, we detect downregulation of Adamtsl2 and Htra3 gene expression in vascular endothelial cells of the hemorrhagic brain. Adamtsl2 is a secreted glycoprotein that regulates ECM assembly necessary in NVUs. Mutations in Adamtls2 that diminish its expression and functions have been linked to connective tissue abnormalities and skeletal dysplasia ^30^ suggesting a key role in normal development. Htra3 is a secreted serine protease which promotes angiogenic sprouting and tip cell formation through mediating endothelial cell-ECM interactions A knockdown of^31^. ^31^. Reduced Htra3 expression in endothelial cells results in decreased developmental angiogenesis ^31^. ^31^. Given that Adamtsl2 and Htra3 are linked to TGFβ signaling, ^32,33^ we predict that dysregulated integrin-mediated TGFβ activation contributes to weakened endothelial barrier integrity. SNPs in the human ITGB8 gene that alter integrin protein expression and TGFβ activation ^34^ have been identified in patients with brain vascular malformations ^35^ and ICH ^36^. Some skeletal dysplasia patients with mutations in ADAMTSL2 have coincident brain vascular pathologies . Interestingly, previous reports have shown that a coordinated expression of Adamtsl2, Htra3, and Spock2 are required for proper Wnt/β-catenin signaling necessary for vascular stability, with a downregulation of these genes leading to impaired BBB development ^37^.^37^. These results support cross-talk between canonical Wnt and TGFβ signaling pathways in NVU development, which we have previously reported ^17^.

Our results also show a significant downregulation of Spock2 expression in endothelial cells of hemorrhagic brains. Downregulation of Spock2 expression disrupts matrix metalloproteinase-2 (MMP2) regulation, driving the degradation of ECM components and activation of integrin signaling pathways ^38^.^38^. This unregulated MMP activity may lead to ECM degradation and leaky blood vessels contributing to endothelial cell proliferation and survival during developmental angiogenesis.**Error! Bookmark not defined.** Nid1 is another ECM r component of the vascular basement membrane, linking laminins, collagens, and proteoglycans to cell surface receptors ^39^. Nid1 is another ECM protein and major component of the vascular basement membrane, linking laminins, collagens, and proteoglycans to cell surface receptors ^39^. Considering the role that these ECM proteins play in basement membrane organization and stability, we predict their decreased expression leads to diminished adhesion of endothelial cells to the ECM, resulting in impaired vascular integrity and subsequent ICH.

At later stages of development during ICH resolution (P5-P10) we observed the upregulation of several genes which point to BBB repair and vascular cell remodeling. Specifically, expression of Ets2, Fam13c, Lrrc32, Ccdc85a, Gabarapl1, and Net1 was upregulated in ICH brains at P5 and P10. Notably, GO enrichment analysis recapitulated this time-dependent pattern of endothelial cell activity in response to ICH. Indeed, while P0 ICH samples show disrupted endothelial cell-ECM adhesion and vascular damage, P5 and P10 endothelial cells show signs of direct response to oxidative stress (response to iron and response to hypoxia) and vascular repair (wound healing, endothelial cell development, and blood vessel remodeling). Notably, the transmembrane protein Lrrc32/Garp, which coordinates with β8 integrin to activate latent-TGFβ ^40^, was upregulated at P5 and, P10. This upregulation of Lrrc32 may represent a compensatory mechanism regulating TGFβ activity and restoring normal blood vessel functions. Considering that TGFβ signaling is necessary for normal microglia function and development ^41,42^, elevated levels of Lrrc32 may restore proper microglia maturation. Notably, Timp3, an MMP inhibitor, was upregulated in endothelial cells at all ages. Given that MMPs play a key role in degrading the ECM, upregulated Timp3 suggests a compensatory mechanism in response to ECM degradation and ICH.

The Gas6/Axl signaling axis is particularly important for modulating the inflammatory response during hemorrhage repair, with peak activity occurring during the subacute phase of injury when iron-handling and inflammatory modulation are most critical ^43^. When activated, Axl suppresses pro-inflammatory cytokine production through inhibition of Toll-like receptor signaling and NF-κB pathways in microglia and macrophages. This anti-inflammatory effect is thought to help prevent excessive inflammation while promoting a more controlled healing response and the survival of surrounding neural cells during the repair process through activation of PI3K/Akt and ERK pathways ^44^, which protect against oxidative stress-induced apoptosis.^44^, which protect against oxidative stress-induced apoptosis. Interestingly, we observed a robust microgliosis in ICH samples, and the single cell transcript data revealed a neuroinflammatory response activated in part by the Gas6-Axl signaling axis in microglial cells. We propose that the upregulation of these components is a key step in ICH clearance and BBB repair. Gas6/Axl signaling influences the expression of Hmox1 and Slc40a1 ^45^, key proteins involved in heme degradation and iron export which we also found to be differentially regulated in microglia. Understanding these mechanisms has potential therapeutic implications, as targeting the Gas6/Axl pathway could provide new approaches for treating hemorrhagic stroke.

Following hemorrhage, the breakdown of hemoglobin from extravasated blood results in a highly toxic iron overload in the brain parenchyma that can lead to neural cell dysfunction and death via ferroptosis . Our results highlight key genes upregulated in microglia which may be critical for the prevention of iron overload and ICH clearance. We propose that Hmox1, Cp, and Slc40a1 work in concert with transferrin to maintain proper iron transport and restore homeostasis after ICH. Previous studies have shown that Hmox1 activity is necessary to attenuate cell death and mediate blood clearance after subarachnoid hemorrhage ^46^. Cp is crucial for efficient hematoma resolution, and Slc40a1 expression is significantly upregulated after ICH as a protective mechanism to export excess iron and prevent iron-mediated oxidative damage. This coordinated system of Hmox1, Cp, and Slc40a1 is particularly important in cells that store or process large amounts of iron, such as macrophages ^47^. Disruption of any component of this system can lead to iron handling disorders; for example, aceruloplasminemia, a rare genetic disorder characterized by Cp deficiency, leads to progressive neurodegeneration due to iron accumulation in the brain. Cp-/- mice develop age-dependent iron accumulation in various brain regions, highlighting its critical role in preventing iron-mediated toxicity ^48^.

Understanding the molecular mechanisms that drive ICH initiation and repair has important therapeutic implications. From a preventive standpoint, maintaining proper integrin-mediated TGFβ activities could protect against spontaneous ICH, particularly in individuals with genetic predisposition to cerebrovascular disorders. In terms of treatment, elucidating the mechanisms that facilitate hemorrhage resolution could reveal new therapeutic targets. These might include alternative pathways for TGFβ activation or downstream effectors that promote vascular repair and stability. We propose the coordinated function of Timp3 regulating ECM stability and iron handling mechanisms (e.g., Hmox1, Cp, and Slc40a1) promoting iron degradation and transport result in hemorrhage clearance. Future research focusing on the temporal dynamics of ICH resolution, combined with detailed molecular analysis of compensatory pathways, may uncover novel therapeutic approaches for treating ICH and other cerebrovascular disorders.

## Materials and Methods

### Experimental mice

Animals used in this experiment were bred in the McCarty lab at the University of Texas MD Anderson Cancer Center and were cared for following IACUC approved breeding and husbandry protocols. Itgb8 mutant mouse models were generated following the breeding schema previously described ^49^.^49^. All mouse tissue samples were collected following IACUC approved protocols and in accordance with ethics and regulations of The University of Texas MD Anderson Cancer Center. Samples include mutant (KO) Nestin-Cre;Itgb8f/f conditional knockout (n=2) and control (WT) Nestin-Cre;Itgb8 +/+ (n=3) mice, collected across three developmental ages (P0, P5 and P10).

### Single-cell RNA sequencing and spatial transcriptome profiling

ScRNAseq was performed on isolated cerebral cortices of two mutant mice and one wild-type control mouse (used twice) for three ages (P0, P5, and P10). Following cortex dissection, formaldehyde fixed, and paraffin embedded (FFPE) neural tissue was dissociated for Chromium Next GEM Single Cell Fixed RNA Profiling following manufacturer protocol (CG000553; 10X Genomics), using manufacturer profiling reagent kit (CG00527; 10X Genomics) and manufacturer Chromium mouse transcriptome probe set v1.0.1.

Xenium *in situ* spatial transcriptomics was performed on sagittal sections of FFPE embedded neural tissue from one mutant and one control mouse from our colony. Samples were processed and analyzed following manufacturer-designed protocol for Xenium Prime with optional Cell Segmentation (User Guide: CG000760, 10X Genomics, Pleasanton, CA).

Fluorophore-tagged probes used in our Xenium experiment included the Xenium Prime 5K Mouse Pan Tissue and Pathway Panel as well as a 100 gene custom probe set containing genes of interest following the scRNAseq data.

### Data processing and quality control

ScRNAseq data were processed using Cell Ranger (v5.0) from 10x Genomics for demultiplexing, alignment, and gene expression quantification. To further remove background counts from ambient RNA and barcode swapping, CellBender (v0.3.0) was applied to the gene-barcode matrices. Although whole brain samples were used in our Xenium data, only cortex cell IDs were extracted from the samples using the Explorer software (10X Genomics) and used for downstream analysis. Downstream analysis of scRNAseq and Xenium data was performed using the Seurat R package (v4.2.1) on RStudio 4.4.1. Cells were filtered based on quality control metrics, including the number of detected genes, total RNA counts, and mitochondrial gene content. After normalization and scaling, samples were integrated using Seurat’ s integration workflow (scRNAseq data) and harmony integration (Xenium data) to correct for batch effects and enable direct comparison between conditions. Dimensionality reduction was performed using principal component analysis (PCA), followed by uniform manifold approximation and projection (UMAP) for visualization. Clusters were identified with graph-based clustering, and conserved marker genes were identified across conditions. Canonical marker genes were used to assign cell types. Bar plots were used to visualize percentage of cell types in each sample. Dot plots were used to visualize transcript percent expression and average expression of gene markers per cell type.

### Differential gene expression and GO enrichment analysis

Differentially expressed genes (DEGs) between conditions were determined using the Wilcoxon rank-sum test from the FindAllMarkers and FindMarkers functions in the Seurat workflow. Unless stated otherwise, a log2fold-change (avg_log2FC) greater than 1 or less than −1 and an adjusted p-value (Bonferroni correction for multiple comparisons) less than 0.05 were used for cutoffs of statistical significance. Differential gene expression comparisons were performed across conditions for each age as well as across ages within conditions. Gene Ontology analysis was performed using the clusterProfiler and org.Mm.eg.db packages to identify biological processes overrepresented in our differential gene expression analysis. The pairwise_termsis function from the enrichplot package was used to construct treeplots which summarize enrichments results by calculating semantic similarity between GO terms by implementing a Jaccard correlation coefficient.

### Slingshot pseudotime trajectory analysis

All integrated Seurat objects previously annotated and used for scRNAseq differential gene expression analysis were merged for pseudotime trajectory inference using the slingshot package. From this merged Seurat object, individual cell types were subset and re-clustered for separate analysis. Using dimensionality reduction clusters from Seurat, the slingshot package creates a minimum spanning tree to identify a lineage structure within the samples and creates a smooth representation of each lineage using simultaneous principal curves. Trajectory analysis was performed separately for each condition. Seurat clusters within trajectories were subset for downstream differential gene expression analysis and GO enrichment analysis.

### Immunofluorescent staining

Brains were harvested and fixed overnight in cold 4% paraformaldehyde (PFA) in PBS. Fixed brains were washed three times in PBS and embedded in paraffin. Formalin-fixed, paraffin-embedded brain tissues were deparaffinized at 65C and rehydrated in decreasing concentrations of alcohol. Heat-induced antigen retrieval was performed in the presence of TE buffer (10mM Tris, 1mM EDTA, pH=9).Sections were permeabilized with 0.2% Triton X-100 in PBS and blocked in 10% normal horse serum for 1 h at room temperature. Tissues were then incubated with the primary antibodies diluted in 1% normal donkey serum, 0.1% BSA, and 0.3 Triton x-100. Primary antibodies were as follow: goat-anti CD31 (AF3628; 1:200), chicken anti-GFAP (05198; 1:1000, Novus), and rabbit anti-Iba-1 (019-19741; 1:200). After washing with PBS, sections were incubated for 45 minutes at room temperature with fluorochrome-conjugated secondary antibodies as follow: anti-rabbit-Alexa Fluor 488 (1:500; 711-545-152, Jackson ImmunoResearch), anti-chicken-Alexa Fluor 647 (1:500; 703-605-155, Jackson ImmunoResearch), and anti-goat-Alexa Fluor 594 (1:500; 705-585-147, Jackson ImmunoResearch).

### Immunohistochemistry staining

Brains were harvested and fixed overnight in cold 4% paraformaldehyde (PFA) in PBS. Fixed brains were washed three times in PBS and embedded in paraffin. Formalin-fixed, paraffin-embedded brain tissues were deparaffinized at 65C and rehydrated in decreasing concentrations of alcohol. Heat-induced antigen retrieval was performed in the presence of TE buffer (10mM Tris, 1mM EDTA, pH=9) for 8 minutes. Sections were blocked in 10% normal donkey serum for 1 h at room temperature. Tissues were then incubated overnight at 4C with the primary antibodies diluted in 10% normal donkey serum. Primary antibodies were as follow: rabbit anti-GFAP (Z0334; 1:500, Dako), and rabbit anti-Iba-1 (013-27691; 1:250, Wako). After washing three times for 10 min. with PBS, sections were incubated for 1 hour at room temperature with secondary antibody horse anti-rabbit (1:500, BA-001, Vector Laboratories). Followed by three washes in PBS for 15 minutes and incubated for 45 minutes in Vector Laboratories ABC kit solution (Vectastain Elite ABC Kit, PK-6100). Slides were washed three times for 15 minutes in PBS and developed in ImmPACT DAB Substrate Kit solution (SK-4105, Vector Laboratories) until background becomes evident (∼5 min) and imaged.

### CellChat Analysis

CellChat v2.1.0 was used to analyze region-specific cell-cell interactions and identify key signaling pathways along with their corresponding ligand-receptor pairs. The analysis was conducted in a contact- or spatial distance-dependent manner, utilizing the truncated mean method to compute average gene expression and communication probabilities of signaling pathways. To identify differences in cell–cell communication strength between WT and KO groups across developmental timepoints (P0, P5, P10), we performed a differential interaction analysis using the Wilcoxon rank-sum test. For each signaling pathway and timepoint, we grouped communication probabilities (prob) by condition (WT vs. KO) and tested for significant differences using rstatix::wilcox_test. Only ligand–receptor (LR) interactions present in both conditions were included, and any missing or zero-probability interactions were excluded. The resulting p-values were adjusted for multiple testing using the Benjamini–Hochberg method to control the false discovery rate (FDR). Significance was assigned as follows: p < 0.05 (*), p < 0.01 (**), and p < 0.001 (***).

### Data access statement

All scRNAseq and spatial transcriptomic data will be deposited in the NCBI GEO database.

---

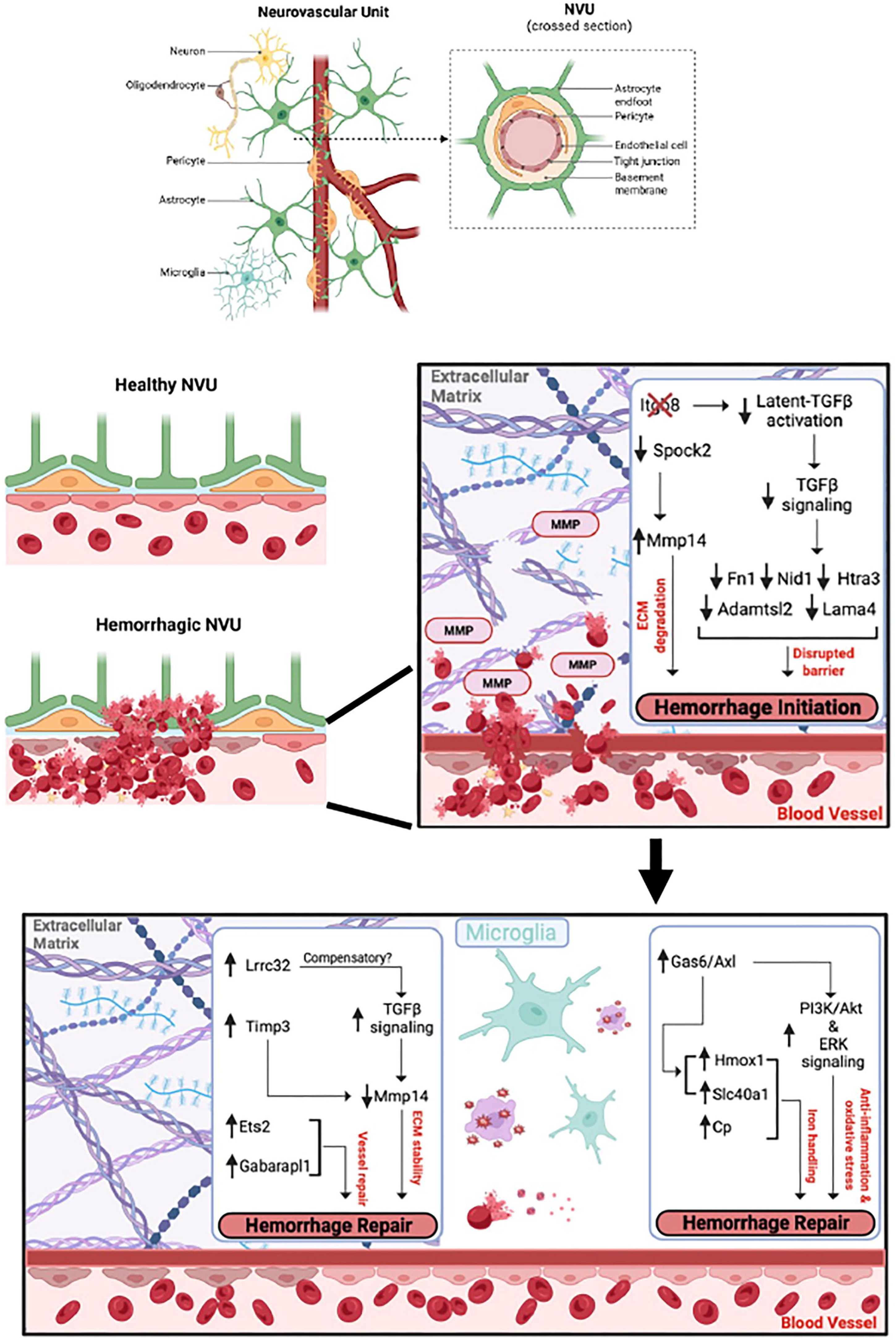

