## Supplemental Materials for "Single Cell RNA Sequencing and Spatial Profiling Identify Mechanisms of Neonatal Brain Hemorrhage Development and Resolution"

**Supplemental Figure Legends**

**Supplemental Figure 1. Reactive gliosis in the hemorrhagic neonatal mouse brain. (A);** Sagittal sections through P5 WT (left) and KO (right) cerebral cortices were immunohistochemically labeled with anti-Iba1 (upper panels) or anti-GFAP (lower panels). Note the increased numbers of reactive Iba1$^+$ microglial cells and GFAP$^+$ astroglial cells in the hemorrhagic cortex (arrows). **(B);** Sagittal sections through the P10 WT (left) or KO (right) cerebral cortex were immunostained with anti-Iba1 (upper panels) or anti-GFAP (lower panels). Note the reactive Iba1$^+$ microglial cells and GFAP$^+$ astroglial cells in the KO sample (arrows).

**Supplemental Figure 2. Analysis of Itgb8 mRNA expression levels in single cells. (A);** Dot plot of Itgb8 expression in astrocytes for each individual P0 KO and WT sample (top left). Feature Plot of Itgb8 expression in astrocyte clusters for combined P0 KO and WT samples (top right). Dot Plot of Itgb8 expression in all cell types for P0 KO and WT samples. **(B);** Dot plot of Itgb8 expression in astrocytes for each individual P5 KO and WT sample (top left). Feature Plot of Itgb8 expression in astrocyte clusters for combined P5 KO and WT samples (top right). Dot Plot of Itgb8 expression in all cell types for P5 KO and WT samples. **(C);** Dot plot of Itgb8 expression in astrocytes for each individual P10 KO and WT sample (top left). Feature Plot of Itgb8 expression in astrocyte clusters for combined P10 KO and WT samples (top right). Dot Plot of Itgb8 expression in all cell types for P10 KO and WT samples.

**Supplemental Figure 3. GO enrichment analysis of DEGs in endothelial cells and microglia. (A);** TreePlot of top 15 gene ontology (GO) enrichment pathways expressed at lower levels in endothelial cells from KO mice at P0 (top), P5 (middle) and P10 (bottom). **(B);** TreePlot of top 15 gene ontology (GO) enrichment pathways activated at lower levels in microglia from KO mice at P0 (top), P5 (middle) and P10 (bottom).

**Supplemental Figure 4. Immunofluorescence analysis of Hmox1 expression. (A, B);** Sagittal sections through the cerebral cortices of P0 and P5 WT (A) and KO mice (B) were stained with H&E (upper panels) or immunofluorescently labeled with anti-CD31 (red) and anti-Hmox1 (green) antibodies. Note the upregulation of Hmox1 protein expression in perivascular microglial cells in control and hemorrhagic cortices at P0 and P5 (arrows). Scale bars, 100 μm.

**Supplemental Figure 5. CellChat analysis of cortical cell-cell communication pathways from Xenium spatial profiling.** Heatmap of cell–cell communication probabilities between sender and receiver cells (y-axis) across all signaling pathways for all cell types (x-axis).

**Supplemental Figure 6. CellChat analysis of cortical cell-cell communication pathways from Xenium spatial profiling. (A-C);** Volcano plots showing log2FC of cell-cell communication probabilities for endothelial cells (A), astrocytes (B), and microglia (C) cells. Significant pathways differentially active in ICH versus control brain samples are displayed by their significance level (ns – grey, $p < 0.05$ - blue, $p < 0.01$ - orange, $p < 0.001$ - red).

Supp. Figure 1

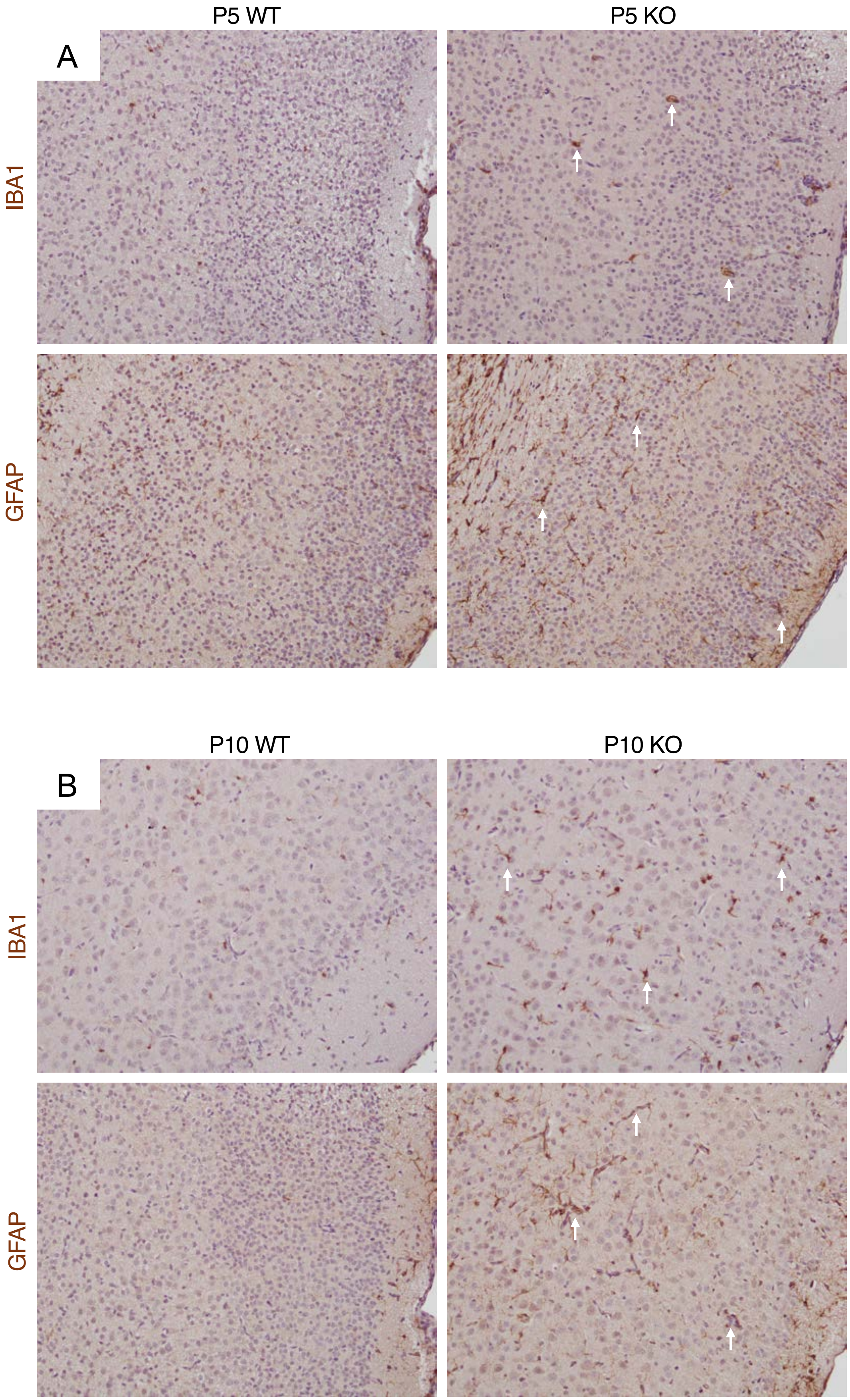

Supp. Figure 2

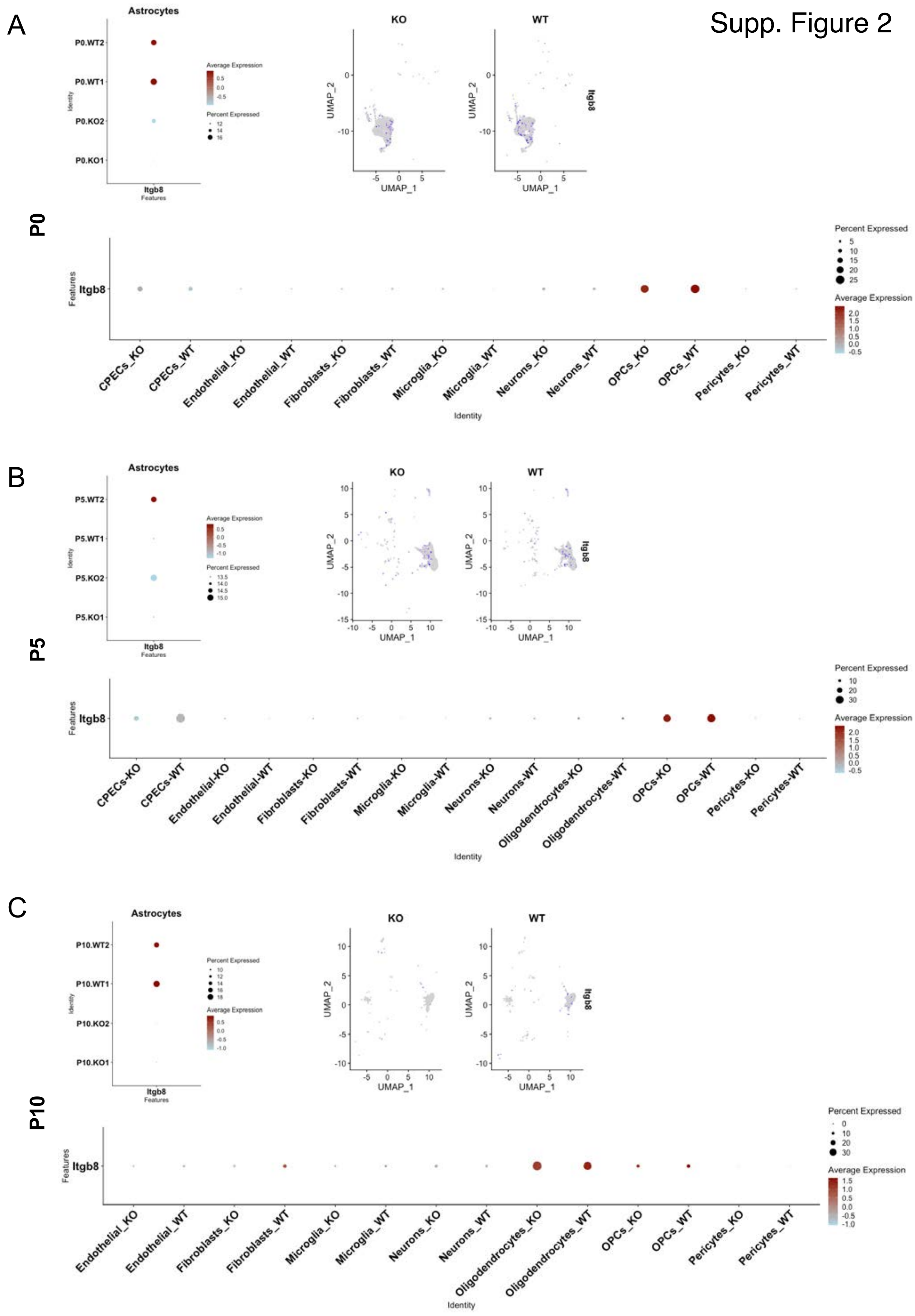

Supp. Figure 3

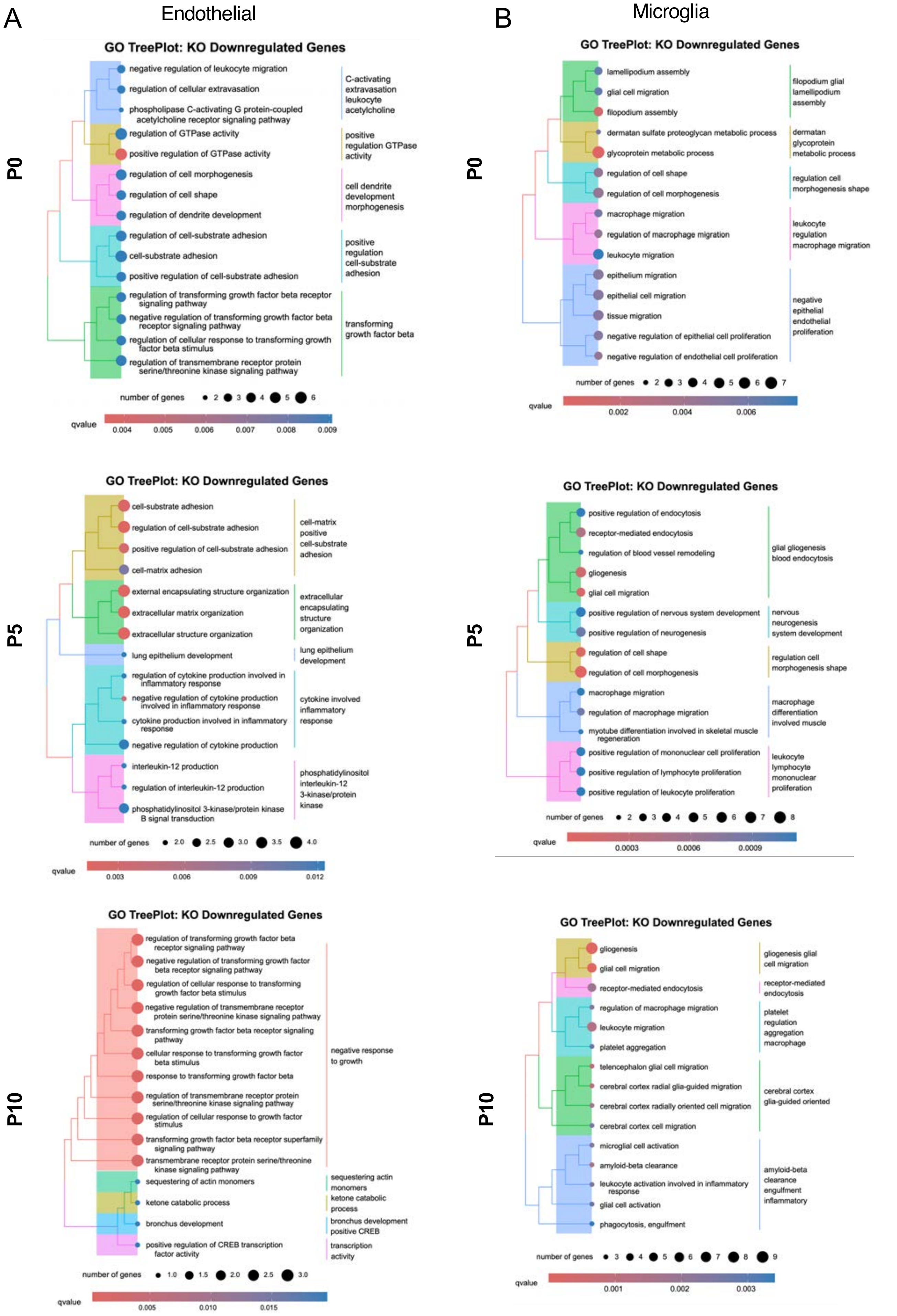

Supp. Figure 4

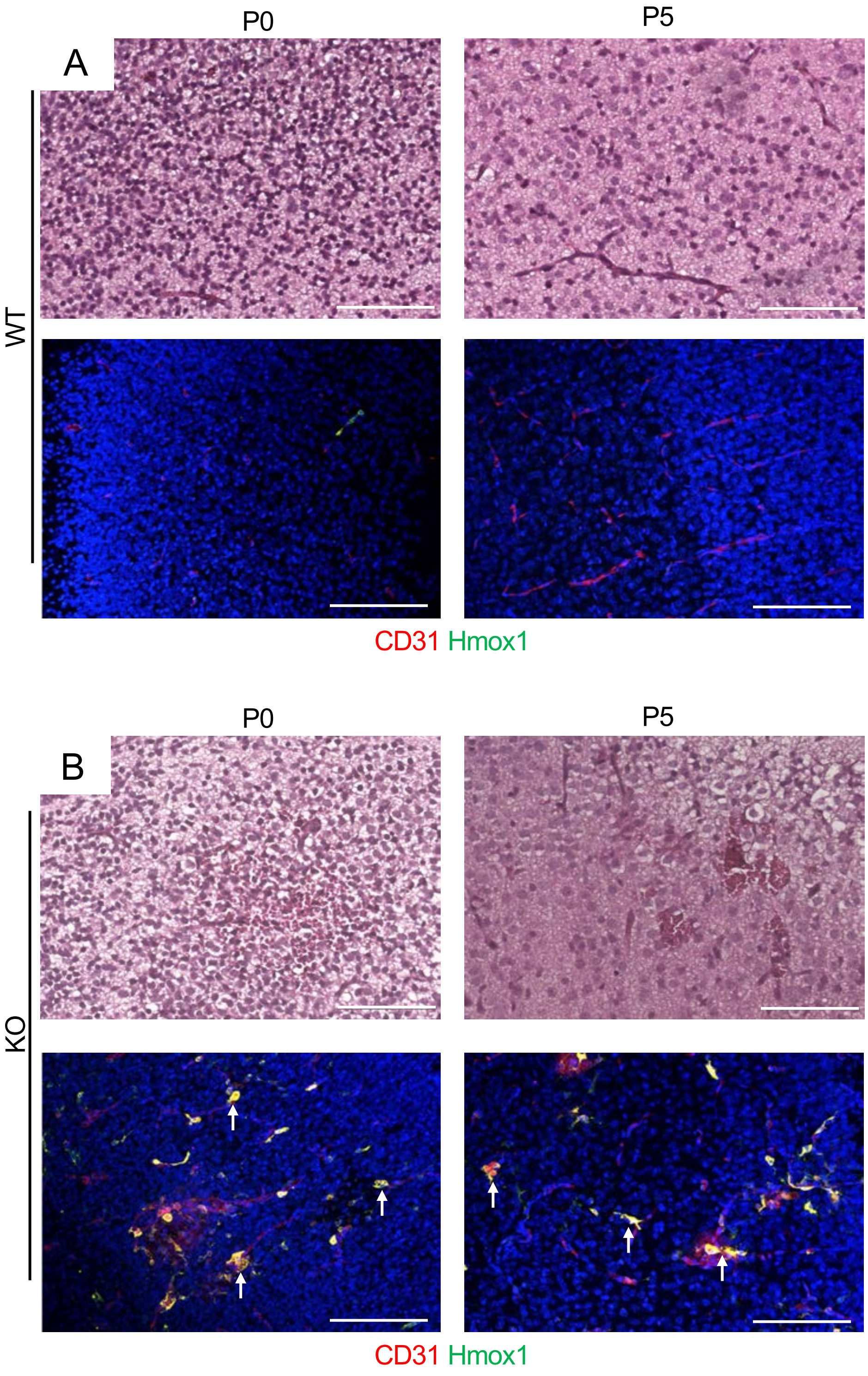

Supp. Figure 5

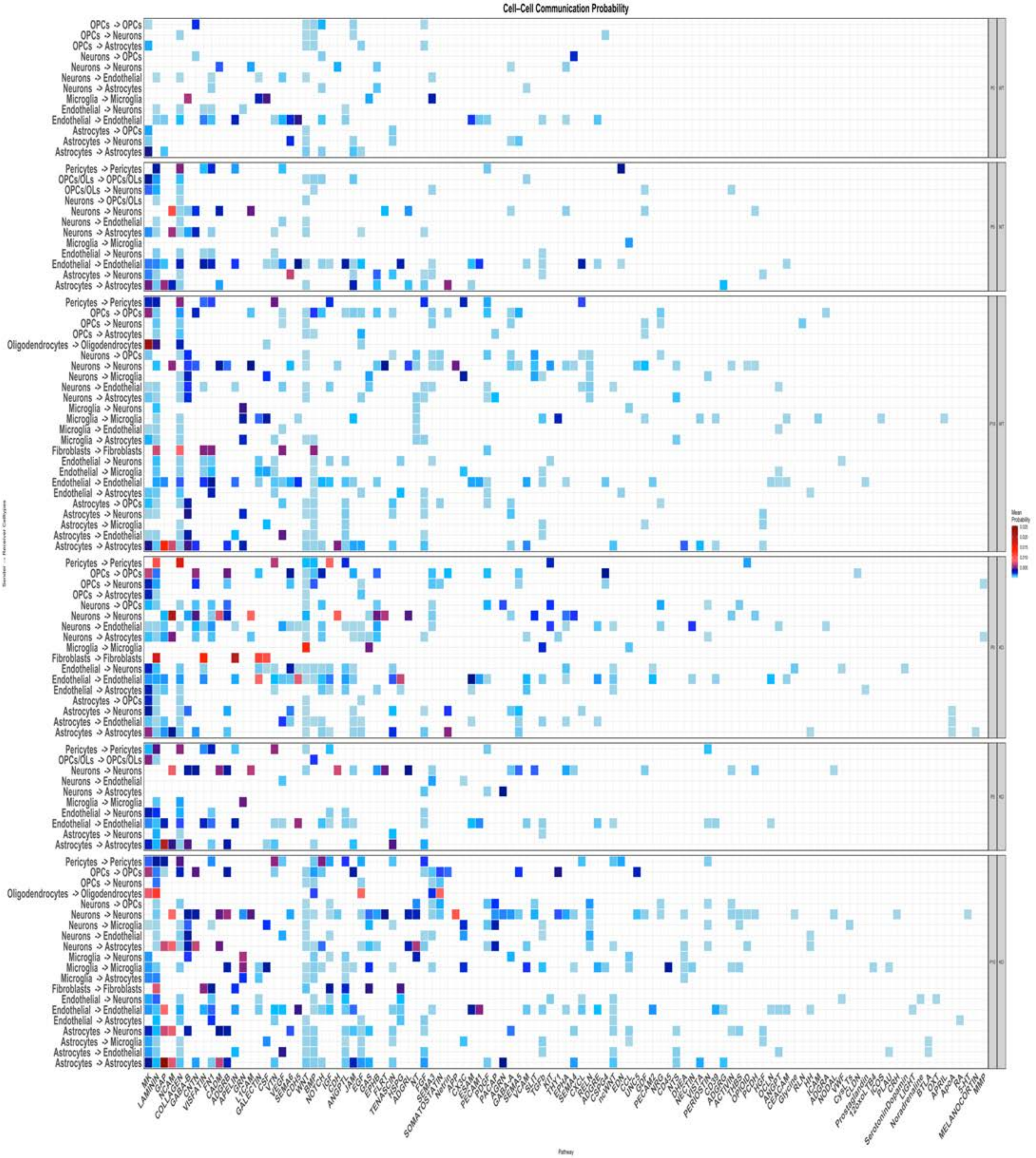

Supp. Figure 6

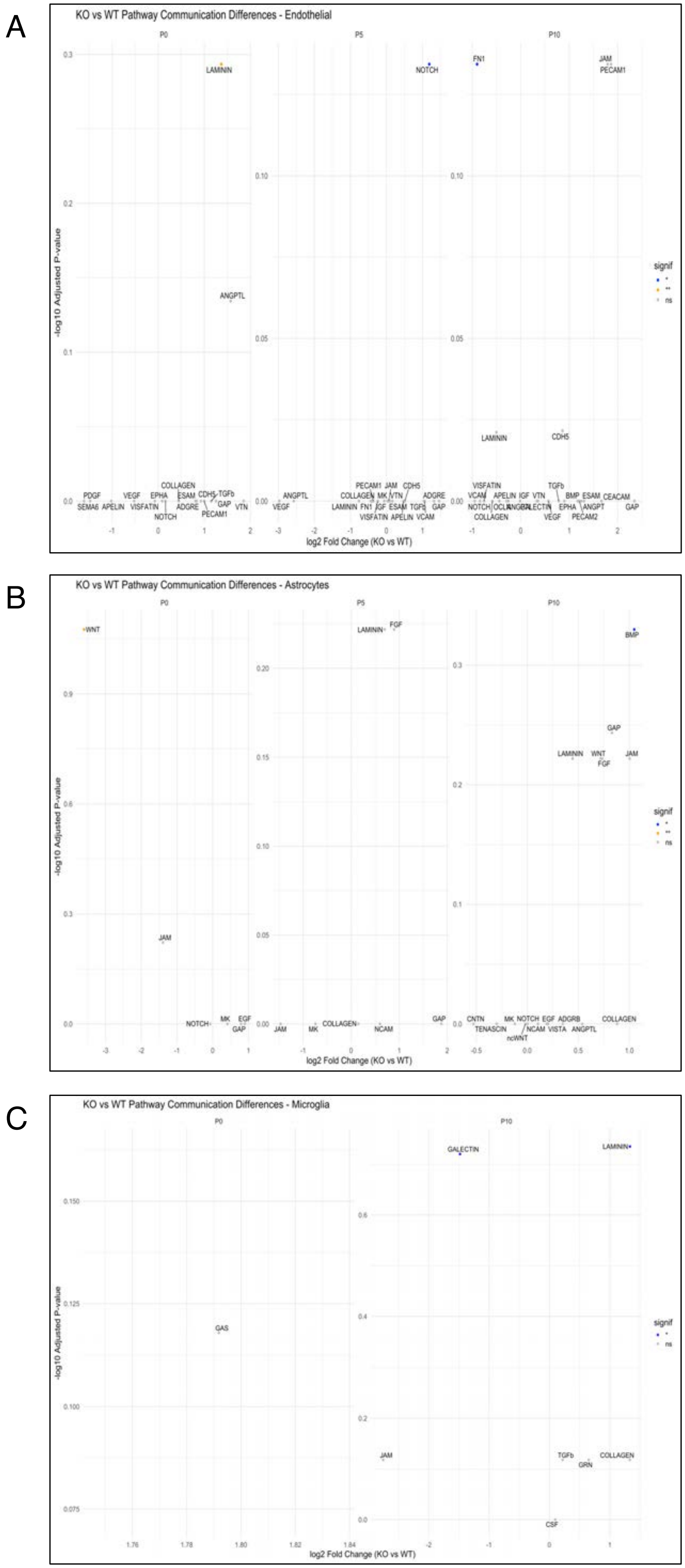
